# KsgA facilitates ribosomal small subunit maturation by proofreading a key structural lesion

**DOI:** 10.1101/2022.07.13.499473

**Authors:** Jingyu Sun, Laurel F. Kinman, Dushyant Jahagirdar, Joaquin Ortega, Joseph H. Davis

## Abstract

Ribosome assembly is orchestrated by many assembly factors, including ribosomal RNA methyltransferases whose precise role is poorly understood. Here, we leverage the power of cryo-EM and machine learning to discover that the bacterial methyltransferase KsgA performs a novel “*proofreading*” function in assembly of the ribosomal small subunit by recognizing and partially disassembling particles that have matured but are not competent for translation. We propose that this activity allows inactive particles an opportunity to reassemble into an active state, thereby increasing overall assembly fidelity. Detailed structural quantifications in our datasets additionally enabled expansion of the Nomura assembly map to highlight rRNA helix and r-protein interdependencies, which newly details how binding and docking of these elements are tightly coupled. These results have wide-ranging implications in our understanding of the quality control mechanisms governing ribosome biogenesis, and showcase the power of heterogeneity analysis in cryo-EM to unveil functionally relevant information in biological systems.

## INTRODUCTION

During ribosome biogenesis in *Escherichia coli*, three ribosomal RNAs(rRNAs) and 54 proteins(r-proteins) assemble into discrete small (30S), and large (50S) subunits that later associate to form a functional 70S ribosome (Shajani et al., 2011). Throughout assembly, r-proteins aid rRNA folding by binding and stabilizing transient RNA folding states (Duss et al., 2019; Rodgers and Woodson, 2019). Concomitant with these folding events, at least 22 known methyltransferases site-specifically modify the rRNA, resulting in 10 and 14 rRNA methylation marks on the 30S and 50S subunits, respectively (Machnicka et al., 2013; Popova and Williamson, 2014). The precise impact of many of these marks on ribosome assembly and function remains unclear (Pletnev et al., 2020).

Whereas the conservation of rRNA methylation sites is generally low between prokaryotes and eukaryotes, two adjacent adenosines (A1518 and A1519; *E. coli* numbering) located in helix 45 are notable for their conservation across the three kingdoms of life (Brimacombe, 1995; Van Knippenberg et al., 1984). Dimethylation at the N6 position of these adenosines is catalyzed by the universally conserved, but not universally essential KsgA/Dim1p enzyme family (Lafontaine et al., 1998; Poldermans et al., 1979a; Poldermans et al., 1979b; Poldermans et al., 1979c).

Though these rRNA methylations are not essential for survival of *E. coli* in laboratory conditions, KsgA homologs confer significant fitness advantages under stress conditions in many organisms. In *Staphylococcus aureus*, for example, KsgA-dependent rRNA methylation increases virulence and contributes to cell survival under oxidative conditions (Kyuma et al., 2015). Similarly, KsgA-deficiency in *Salmonella enterica* confers susceptibility to high osmolarity and attenuates virulence (Chiok et al., 2013). Early studies established the general belief that these phenotypes were the consequence of KsgA-dependent methylations fine-tuning the structure of the ribosome and ultimately contributing to its fidelity and overall translation efficiency (Cunningham et al., 1990; Mangat and Brown, 2008). More recent work, however, has identified phenotypes in Δ*ksg*A *E. coli* (Connolly et al., 2008) that are commonly found in strains lacking well-established ribosome biogenesis factors such as YjeQ, RimM, and Era (Himeno et al., 2004; Leong et al., 2013; Thurlow et al., 2016), leading to the hypothesis that KsgA could also actively participate in ribosome assembly.

To explore the mechanisms through which KsgA assists ribosome assembly, we applied cryo-electron microscopy (cryo-EM) and cryoDRGN (Zhong et al., 2021), a newly developed single particle cryo-EM image processing pipeline, to structurally characterize the ensemble of free 30S assembly intermediates that accumulate in a Δ*ksg*A strain of *E. coli*. Treatment of these purified assembly intermediates with KsgA revealed that it specifically targeted inactive 30S particles and induced large-scale structural remodeling in these particles, suggesting that KsgA acts as a quality control factor during ribosome assembly. Additionally, by leveraging cryoDRGN’s ability to extract and quantify a continuum of structures, we uncovered how assembly of individual rRNA helices and r-proteins influence one another. These analyses enabled construction of an “extended” Nomura assembly map (Held et al., 1974; Mizushima and Nomura, 1970) that newly depicts interdependencies between the native docking of rRNA helices and r-protein binding events, helping to further explain how protein-dependent conformational changes in the 16S rRNA facilitate the high degree of cooperativity observed in ribosome biogenesis (Stern et al., 1989).

## RESULTS

### KsgA impacts assembly of small ribosomal subunits

To investigate the role of KsgA in the assembly of ribosomal small subunits (SSU), we first isolated and characterized ribosomal small subunit assembly intermediates that accumulated in Δ*ksg*A cells (Baba et al., 2006) grown at low temperature. Consistent with a previously reported KsgA-dependent defect in ribosome biogenesis (Connolly et al., 2008), these cells contained more unprocessed 17S rRNA, exhibited a distinct sucrose gradient profile, and the 30S_Δ*ksg*A_ particles isolated from them bore an incomplete complement of ribosomal proteins when compared to that isolated from wild-type cells (**Supplementary Figure 1**). Using microscale thermophoresis, we found that these 30S_Δ*ksg*A_ particles were competent to bind to KsgA *in vitro* (**Supplementary Figure 2**), indicating that the 30S_Δ*ksg*A_ particles we isolated had not matured to a stage beyond which KsgA could act.

Next, we used single particle cryo-electron microscopy (cryo-EM) to visualize the 30S_Δ*ksg*A_ particles and to evaluate the effect of adding KsgA to them *in vitro*. The resulting maps exhibited two striking features. First, both maps bore canonical features of immature SSUs, including incomplete or missing density in helix 44, which is strictly required for subunit joining and subsequent translation (Qin et al., 2012; Schuwirth et al., 2005). Second, each map exhibited a high degree of resolution anisotropy in the head, platform, and spur domains, consistent with conformational or compositional heterogeneity in these regions. Surprisingly, in the KsgA-treated samples, the local resolution in these regions was further diminished, consistent with even greater structural variability (**Figure 1A**).

**Figure 1:**
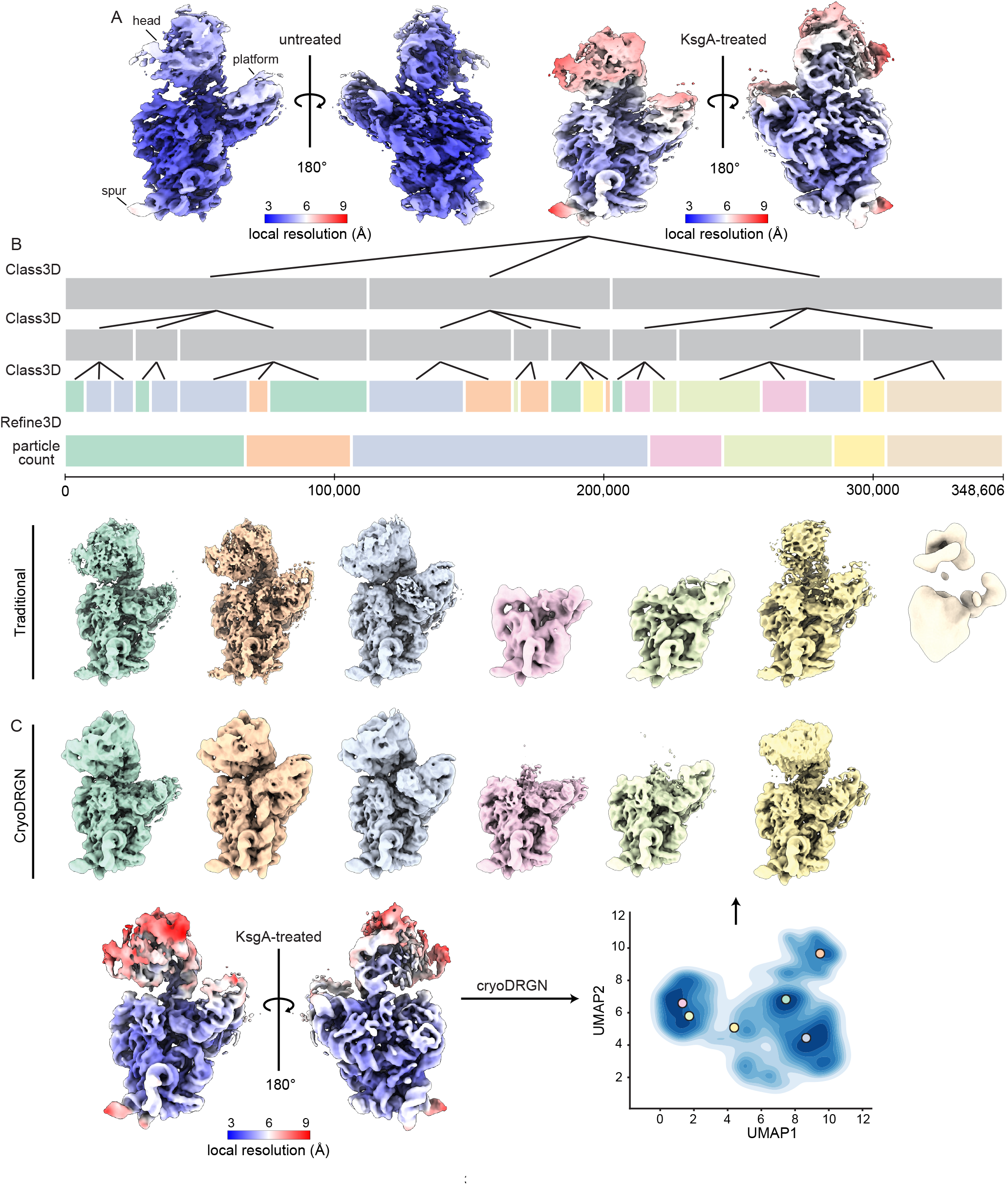
KsgA treatment of 30S_Δ*ksg*A_ particles produces a heterogeneous structural ensemble. (**A**) Density maps of untreated and KsgA-treated 30S_Δ*ksg*A_ particles, colored by local resolution. (**B**) Traditional hierarchical classification and refinement of KsgA-treated particles. Each bar represents a layer of 3D-classification or refinement, with its length proportional to the number of particles in that class. Classes in the final layer are colored to indicate how they were pooled for 3D refinement, and volumes are colored to match. (**C**) Consensus map of KsgA-treated particles included in the final round of cryoDRGN training (see Supplementary Figure 5), colored by local resolution, and UMAP representation of the latent space produced by cryoDRGN analysis of these particles (bottom). Colored markers indicate the position in latent space from which volumes sampled were generated (top).

### KsgA binding remodels large rRNA domains in late-stage 30S assembly intermediates

Given the unexpected increase in structural variability upon addition of KsgA, we aimed to further investigate these structures. Systematically analyzing heterogeneous structural ensembles remains an open challenge in cryo-EM, and thus we employed both RELION’s traditional Bayesian 3D-classification and refinement methods (Scheres, 2016) (**Figure 1B; Supplementary Figures 3**,**4**) and cryoDRGN (Zhong et al., 2021), a neural network-based reconstruction approach that has shown great promise in this arena (Kinman et al., 2022). Specifically, we trained a cryoDRGN neural network model on this dataset, producing a latent encoding for each particle and a trained decoder network capable of producing density maps from any position in latent space supported by underlying particle images (**Supplementary Figure 5A**). Consistent with large-scale, discrete heterogeneity, we found that the latent space was featured, and volumes sampled from various populated locations within the latent space revealed the presence of massive structural variability in the head, platform, and KsgA-binding site (**Figure 1C**). These “major” structural classes were similar to those uncovered using traditional 3D-classification, supporting the accuracy of cryoDRGN’s neural network-based decoder.

To systematically interrogate structural heterogeneity present within the KsgA-treated dataset, we exploited cryoDRGN’s powerful generative model by sampling 500 volumes from the latent space, effectively generating density maps from all regions of latent space that were supported by data (**Supplementary Figure 5B**). To interpret the resulting density maps, we applied a coarse-grained “*subunit occupancy analysis*” (Davis et al., 2016; Davis and Williamson, 2017). Specifically, we measured the amount of density we observed for each rRNA helix and r-protein, relative to that expected based on the atomic model of the fully mature subunit (**Figure 2A**). The resulting heatmap depicted the fraction of natively localized density occupied by each structural element (69 columns) in each density map (500 rows), and revealed that a vast array of structures was present upon addition of KsgA (**Figure 2B**). Using the hierarchically clustered occupancy map, we grouped these structures into classes of broadly similar density maps, and we generated representative structures of each class using cryoDRGN (**Figure 2C, Supplementary Figure 6**, and Methods). Here, we observed both highly mature particles (classes 1-4) and substantially less mature structures (classes 5-11), including maps that lacked density for the head (classes 8-9), platform (classes 5-6), or both (classes 7 and 10).

**Figure 2:**
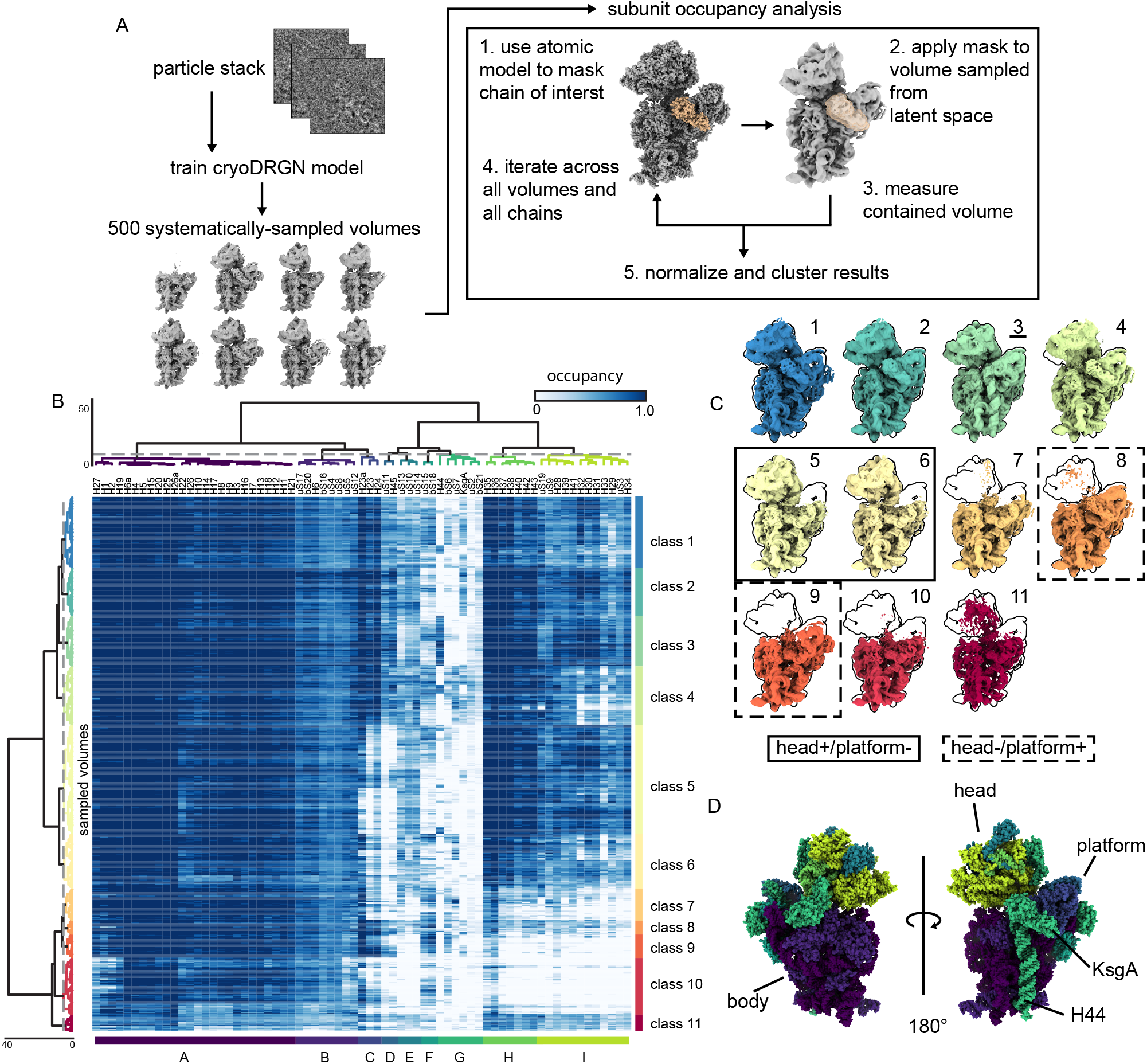
Subunit occupancy analysis of KsgA-treated 30S_Δ*ksg*A_ particles reveals large structural domains that cooperatively interconvert. (**A**) Depiction of our subunit occupancy analysis approach (see Methods). (**B**) Occupancy analysis of KsgA-treated particles displayed as a heatmap. Rows (500) correspond to sampled density maps and columns (69) correspond to structural elements defined by the atomic model. (**C**) Volumes generated from centroid position in latent space of each of the classes shown in (B). Volumes are outlined by the silhouette of the mature 30S (class 3, underlined). Maps with head density but no platform density are highlighted with a solid box; maps with platform density but no head density are surrounded with dashed boxes. (**D**) Atomic models of the 30S subunit used in performing subunit occupancy analysis (PDB: 4V9D, 4ADV) are colored by the structural blocks defined through hierarchical clustering in (B). Structural features of interest are annotated.

### Treatment of 30S_Δ*ksg*A_ particles with KsgA reveals cooperatively interconverting structural blocks

When inspecting the columns of the subunit occupancy heatmap (**Figure 2B**), we noticed structured blocks consistent with the cooperative assembly of the 30S subunit (Davis and Williamson, 2017). The blocks were structurally coherent, containing neighboring rRNA helices and r-proteins, and were largely organized around the known SSU head, platform, and body domains (**Figure 2D**). Careful inspection of the occupancy of structural blocks C/D and H/I, which encompass the platform and head, respectively, revealed that their occupancy was uncoupled, with some volumes bearing only the head, some bearing only the platform, and others having both or neither. In contrast, blocks corresponding to the body (A/B) were always present, consistent with prior work hypothesizing a requirement for formation of the body before assembly of the head or platform (Mulder et al., 2010; Nomura, 1974; Sashital et al., 2014). Overall, these observations support the existence of parallel pathways facilitating the independent assembly of the head and platform domains.

Interestingly, although these structural blocks are largely coherent, consistent with their cooperative maturation, they are not perfectly so. For example, 29 maps in class 4 exhibited low occupancy of helices 32 and 33, which represent the most extreme region of the head domain (**Figure 2B**). Visual inspection of these maps revealed rotations of the head domain away from the body, using the neck that connects the head to the body as a fulcrum. This apparent conformational flexibility likely contributed to the relatively low local resolution of the head density in our traditional 3D reconstructions (**Figure 1B**), and highlighted the power of this cryoDRGN occupancy analysis approach to resolve lowly populated conformers.

### KsgA binds a diverse array of small ribosomal subunits

In addition to systematically enumerating large-scale structural changes, this cryoDRGN-based approach allowed us to thoroughly quantify the presence or absence of proteins and rRNA helices across the dataset. Thus, we could readily determine the fraction of the ribosomal particles bound to KsgA upon treatment. We found that only ∼39% of the particles were bound to KsgA, despite having added the factor super-stoichiometrically and in excess of the apparent K_D_ (**Supplementary Figure 2**). This sub-stoichiometric KsgA occupancy implied that not all of the ribosomal particles were competent to bind KsgA and suggested that KsgA may recognize specific structural features on the ribosome that were not uniformly present across all classes of immature subunits.

To better understand which structural elements KsgA recognizes, we used our subunit occupancy analysis and extracted the subset of maps with high KsgA occupancy. We designated these maps as “KsgA-bound” and performed hierarchical clustering of the occupancy matrix specifically on this KsgA-bound subset of 185 maps (**Figure 3A, Supplementary Figure 6B**). Consistent with recent reports of KsgA bound to a mature 30S subunit (Schedlbauer et al., 2021; Stephan et al., 2021), in these maps, we observed clear density near the decoding center that was well fit by an atomic model of KsgA (O’Farrell et al., 2004) (**Figure 3B-C**). Notably, we found that KsgA bound to particles of highly variable composition, including maps presenting densities for all major domains (body, platform and head; class 3) and maps lacking portions of the platform (class 4) or the head (class 5) (**Figure 3B**). This result highlighted the relative independence of the KsgA binding site and distal elements in the head, and suggested that KsgA is primarily sensing local structural elements on the 30S particles.

**Figure 3:**
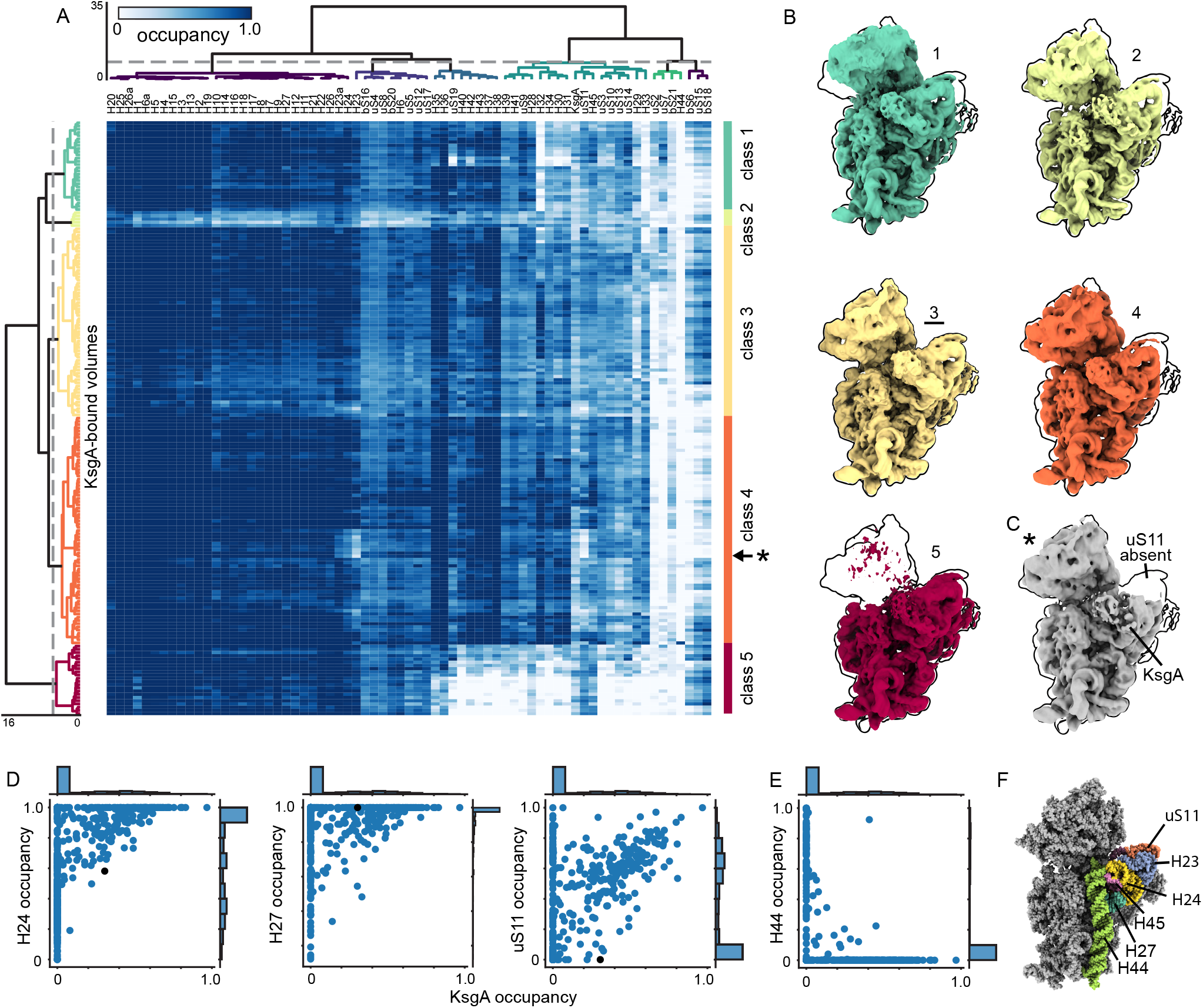
KsgA binds a diverse array of assembly states. (**A**) Re-clustering of KsgA-bound maps (185) based on subunit occupancy, displayed as a heatmap. (**B**) Centroid maps for KsgA-bound classes, outlined by the most mature volume (class 3, underlined). (**C**) A map sampled from row labeled with * in (A). Note platform element uS11 is missing. (**D**) Correlations between occupancy of KsgA and platform elements thought to be critical for KsgA binding. Black dot notes occupancy of volume depicted in (C). (**E**) Correlation between KsgA and H44 occupancy, consistent with mutually exclusive KsgA binding and H44 docking. (**F**) Atomic models of the 30S ribosome (PDB: 4V9D, 4ADV) with structural features annotated. KsgA is hidden to show platform elements.

Detailed inspection of occupancy patterns in ribosomal elements proximal to the KsgA binding site highlighted those dispensable for KsgA association, and those consistently occupied in our structures. Indeed, we found that rRNA helices 24 and 27 are highly occupied in all maps with high KsgA occupancy (**Figure 3D**), consistent with recent results implicating these helices in binding of the C-terminal domain of KsgA (Stephan et al., 2021). This analysis further established the mutually exclusive occupancy of KsgA and helix 44, and it newly highlighted that uS11 in the platform is largely, but not strictly, required for KsgA binding as we identified five KsgA-bound maps lacking uS11 (**Figure 3C-F**).

### RNA backbone contacts facilitate binding of KsgA to the immature 30S_Δ*ksg*A_ particles

To inspect the atomic contacts supporting KsgA binding, we next collected a larger dataset of the KsgA-treated 30S_Δ*ksg*A_ particles and, using hierarchical classification and multi-body refinement in RELION (Nakane et al., 2018; Zivanov et al., 2018), we reconstructed a 2.8 Å resolution map of this complex that grossly resembled that of KsgA bound to mature 30S_Δ*ksg*A_ subunits derived from dissociation of 70S_Δ*ksg*A_ particles (Stephan et al., 2021) (**Figure 4A, Supplementary Figure 7**).

**Figure 4.**
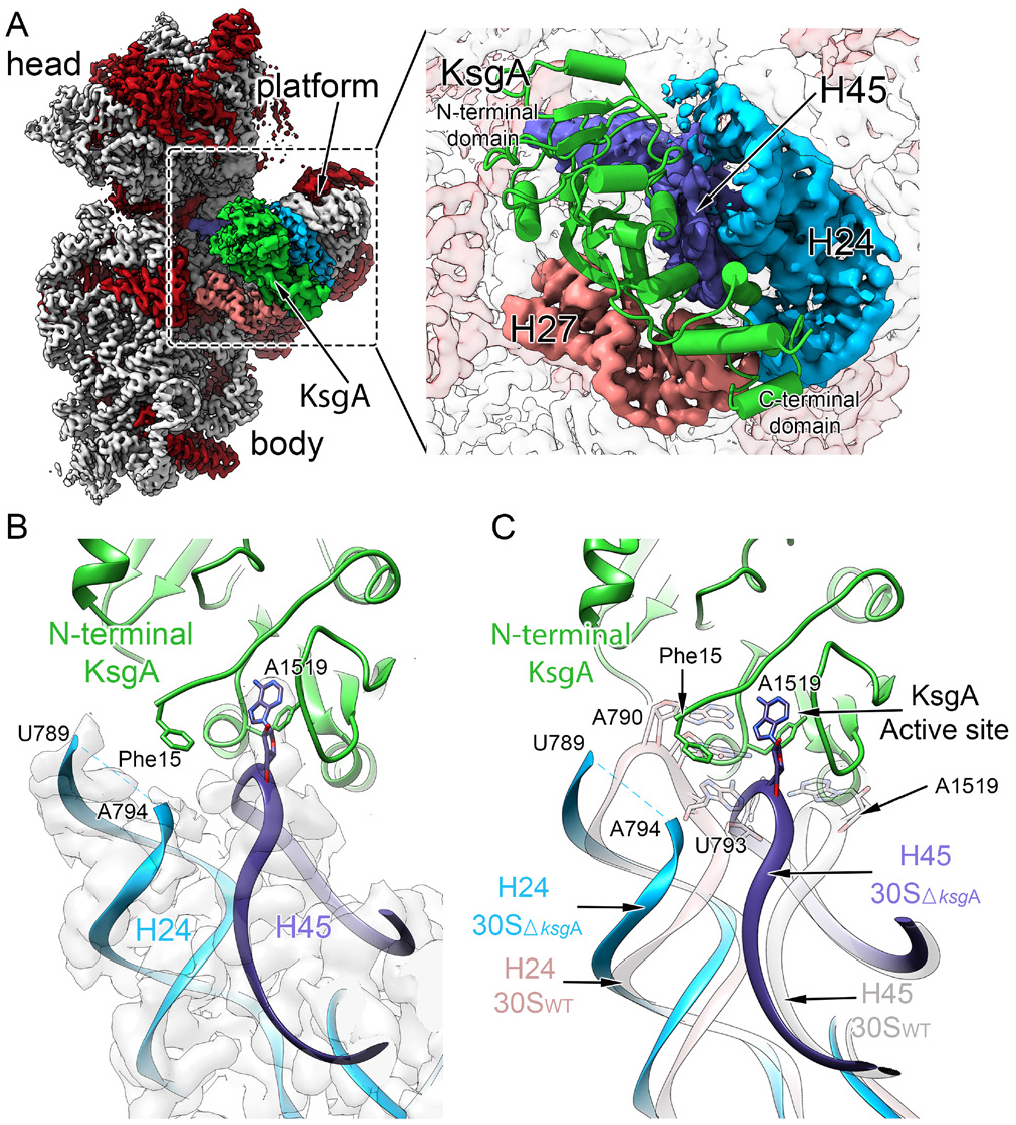
Substrate engagement by KsgA displaces a gatekeeping rRNA helix. (**A**) Interface view of the cryo-EM structure obtained for the immature 30S_Δ*ksg*A_ particle bound to KsgA (green). Ribosomal proteins are shown in red and the 16S rRNA is shown in light gray, and structural landmarks of the ribosomal subunit are indicated. Key rRNA helices interacting with KsgA are colored in pink (helix 27), cyan (helix 24) and blue (helix 45). The interaction area is enlarged in the right panel, and depicts a molecular model of KsgA derived from the cryo-EM map. (**B**) Magnified view of the interface between KsgA’s N-terminal region and rRNA helix 24, and KsgA’s active site and substrate residue A1519 from helix 45. Note the lack of cryo-EM density (gray) corresponding to helix tip residues 790-793, and the proximity of KsgA residue Phe15 to this region. (**C**) Overlay of rRNA helices 24 and 45 from the molecular model of the mature 30S_WT_ subunit in the absence of KsgA (rose) and those from the 30S_Δ*ksg*A_ particle bound to KsgA (H24 in cyan; H45 in purple). Note positioning of A1519 in KsgA’s active site necessitates displacement of helix to avoid steric clashes between helices 24 and 45.

Construction of a molecular model using this map revealed that KsgA binding was primarily supported by contacts to backbone phosphates and sugars of rRNA helices 24, 27, and 45 (**Supplementary Figure 8A-C**), with the substrate rRNA residue (A1519) bound in the KsgA catalytic site and stabilized in this conformation by a π-stacking interaction with KsgA residue Tyr116 (**Supplementary Figure 8D**). In contrast, KsgA substrate residue A1518 was placed away from the active site, suggesting that A1519 is methylated first. We further observed KsgA active site residues Asn113 and Leu114, which are known to facilitate catalysis (O’Farrell et al., 2012), hydrogen bonded to the methyl-receiving N6 atom of A1519, apparently priming it for methylation (**Supplementary Figure 8D**). Notably, this overall positioning of helix 45, which contains the substrate residues, was maintained primarily through KsgA contacts to backbone elements of the rRNA, suggesting that such stabilizing contacts would also be available when A1518 binds in the active site for subsequent KsgA-dependent methylation.

Our model additionally showed that upon binding, KsgA’s N-terminal region approached helix 24 (H24) and we were unable to resolve density corresponding to H24 nucleotides 790-793 (**Figure 4B**), suggesting that KsgA binding induced flexibility in this region. Comparison of our molecular model with that of KsgA-free 30S_WT_ led us to hypothesize that this KsgA-induced remodeling of H24 plays a functional role in regulated catalysis. Indeed, in the canonical conformation, H24 residues 790-793 would protrude into the KsgA active site, with nucleotide U793 precluding A1519 from adopting the “flipped-out” conformation required to access the KsgA active site (**Figure 4C**). As such, we interpreted the coupled H24 and H45 motions upon KsgA binding as catalysis-independent priming of the substrate for methylation.

### Nearly mature 30S subunits accumulate in KsgA’s absence

To understand how the 30S subunit assembles in the absence of KsgA, we applied our cryoDRGN-occupancy analysis pipeline to untreated immature 30S_Δ*ksg*A_ particles (**Supplementary Figure 5, 6C**). Given KsgA’s role as an assembly factor, we expected these structures to generally appear less mature than those observed upon addition of the factor. Instead, we were surprised to find that these ribosomal particles were significantly less heterogeneous than the ones present upon addition of KsgA, with the centroid volumes from occupancy analysis appearing more similar to the mature 30S (**Figure 5A, Supplementary Figure 9**). Indeed, while the head density was weak in one of these ten representative volumes (class 8), only 9 of the 500 sampled volumes, representing only 1% of the full particle stack, lacked head density entirely (**Figure 5B**). By comparison, in the KsgA-treated dataset, 118 of the 500 sampled volumes (classes 7-10, **Figure 2C**), or 20% of the particle stack, lacked head density entirely. To further assess the maturity of untreated versus KsgA-treated particles, we used our 500 sampled volumes from each dataset and calculated a voxel-wise sum of squared residuals between each volume and a paired mature reference volume (see Methods). Plotting this data as a cumulative distribution function (**Figure 5C**) or inspecting density maps sampled at regular intervals along the y-axis of this plot (**Supplementary Figure 10**) highlighted that untreated particles are globally more structurally similar to the mature volume than their KsgA-treated counterparts.

**Figure 5:**
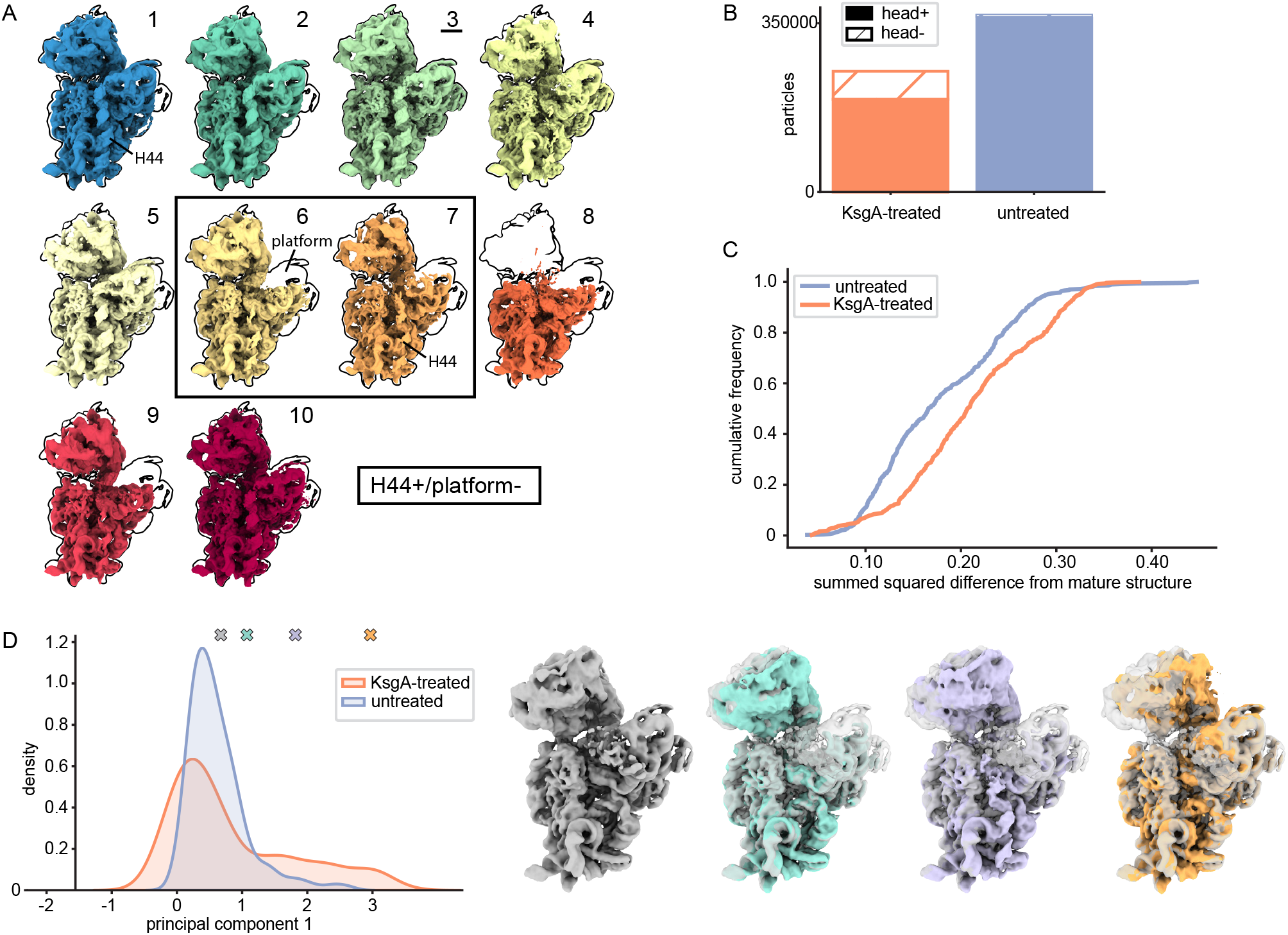
Nearly-mature SSUs accumulate in the absence of KsgA. (**A**) Centroid maps for classes defined by occupancy analysis of untreated particles (see Supplemental Figure 8). Maps are outlined by the most mature class (3, underlined). A solid box surrounds maps that presented helix 44 density but lacked platform density. (**B**) Total number of head+ and head-particles in each dataset. (**C**) Cumulative frequency plot for normalized summed squared difference values calculated between individual binarized maps and the paired mature 30S reference map (see Methods). (**D**) Principal component analysis was performed on the voxels within a mask corresponding to the native H32/H33 region in 500 volumes sampled from the head+ subset of latent space for each dataset. The density distribution along the first principal component is shown for each dataset. Colored markers indicate positions along principal component 1 from which the volumes in the right panel were sampled. The initial gray volume is overlaid with each structure for reference.

### KsgA induces partial particle disassembly

Because particles were less mature upon addition of KsgA, we reasoned that KsgA binding may lead to partial disassembly of nearly-mature 30S_Δ*ksg*A_ particles. To explore this possibility biochemically, we asked whether KsgA binding led to dissociation of r-proteins by treating immature 30S_Δ*ksg*A_ particles with a 10-fold molar excess of KsgA at 37°C for 20 min. Following the incubation, the ribosomal particles were separated from free r-proteins by ultracentrifugation and the pelleted particles were analyzed by quantitative mass spectrometry (Jomaa et al., 2014). Interestingly, we found that the r-protein composition after KsgA-treatment was indistinguishable from that of the untreated immature 30S_Δ*ksg*A_ particles (**Supplementary Figure 1C**), indicating that r-proteins do not appreciably dissociate upon KsgA binding. Instead, these data support a model in which KsgA binding causes uncoupling of the head and body, with each domain remaining folded and bound by r-proteins, but now capable of rotating relative to one another.

To quantify destabilization of the head upon KsgA treatment, we developed a “*voxel principal component analysis*” approach (**Supplementary Figure 11**, and Methods) that leveraged cryoDRGN’s ability to generate many density maps. This approach allowed us to visualize head domain motions within our structural ensembles, and to compare the degree of motion before and after addition of KsgA. Indeed, with this approach, we observed a long-tailed distribution along principal component 1 specifically in the KsgA-treated dataset, and volumes sampled within the tail bore an undocked and rotated head domain (**Figure 5D**). The presence of particles with undocked heads specifically in the KsgA-treated dataset supports a model in which KsgA binding caused uncoupling of the body and head, permitting free rotation of the head and body relative to each other (**Supplementary Video 1**).

### KsgA recognizes the inactive conformation of a key rRNA helix

This apparent KsgA-induced structural uncoupling was surprising and led us to hypothesize that specific particles accumulating in the Δ*ksg*A strain may subtly differ from mature, active 30S_WT_ particles, with this difference allowing for specific recognition and remodeling by KsgA. According to such a “*proofreading*” model (Hopfield, 1974), KsgA would preferentially bind to such particles, induce structural remodeling by uncoupling of the head and body, and thus allow the particle another opportunity to re-assemble into an active form.

To test this hypothesis, we carefully inspected the untreated dataset for evidence of such structures. Helix 44 (H44) is traditionally considered to be one of the last elements of the 30S ribosome to form (Jomaa et al., 2011), and thus we were surprised that the majority (67%) of the 30S_Δ*ksg*A_ particles had significant H44 occupancy and that this proportion decreased to 21% upon addition of KsgA (**Figure 6A**). Visual inspection of the centroid volumes from the untreated sample (**Figure 5A**), and quantitation of H44 occupancy revealed many particles in which H44 was present even in the absence of platform structural elements uS11 and H45, suggesting premature H44 docking (**Figure 6B**). Given these observations and the proximity of H44 to the KsgA binding site, we hypothesized KsgA may be recognizing a structural feature related to this premature H44 docking. In support of this hypothesis, we noted a substantial population of particles displaying H44 occupancy between 0.2 and 0.8, which was suggestive of H44 adopting non-canonical conformations (**Figure 6A**). Indeed, volumes sampled from within this region displayed H44 in an unexpected, but previously reported inactive conformation (Jahagirdar et al., 2020) in which H44 is “unlatched” and moved away from the body of the ribosome. This conformation, which was first discovered in the 1960s by Elson and colleagues (Zamir et al., 1969), is not competent to bind to 50S particles or support translation.

**Figure 6:**
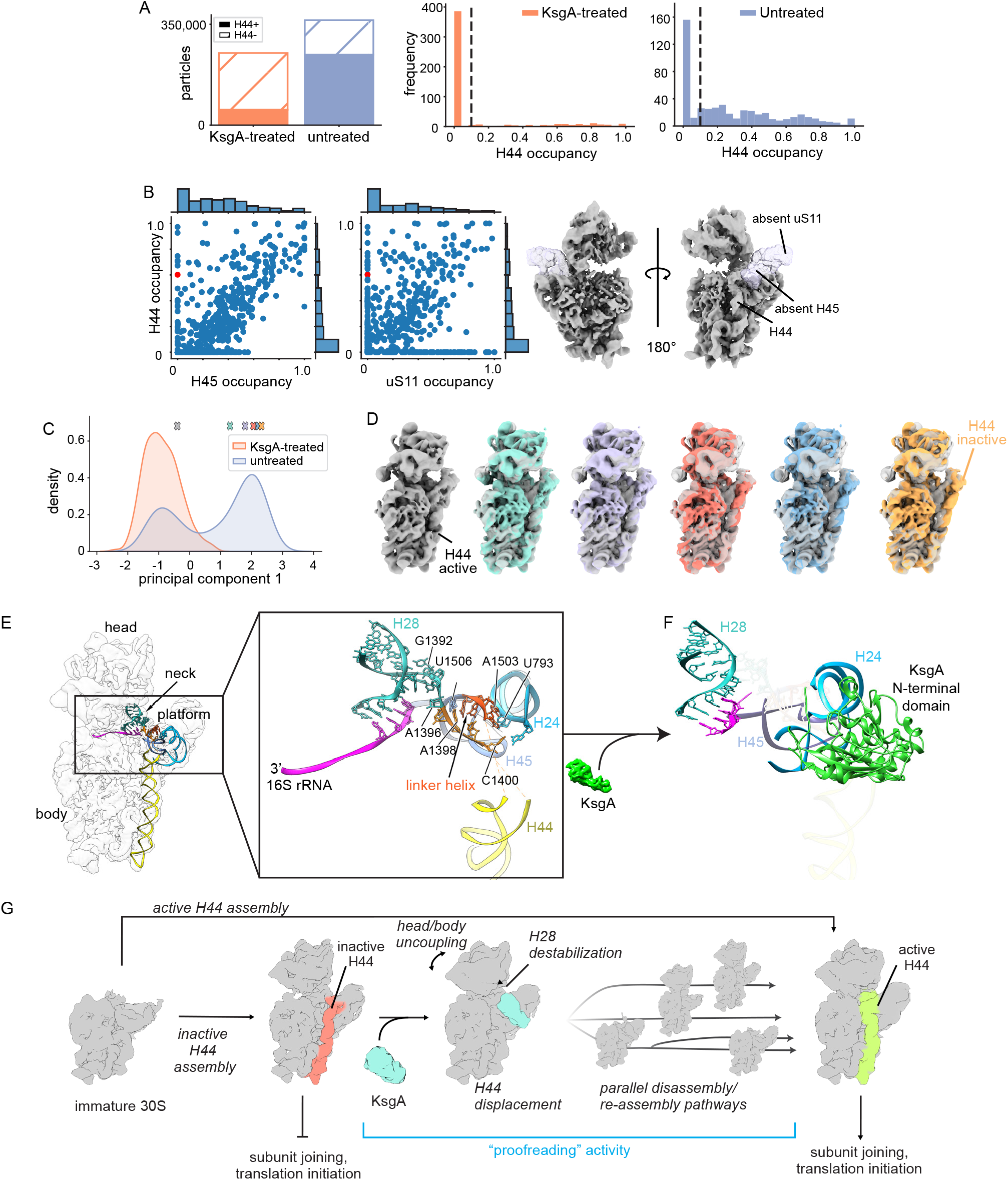
KsgA recognizes and remodels inactive subunits. (**A**) Bar chart displaying the total number of H44- and H44+ particles in each dataset (left), and histograms of the occupancy of helix 44 in the 500 sampled volumes from each of the two datasets (right). The dashed line notes the occupancy threshold we used to distinguish H44+ and H44-particles. (**B**) Occupancy correlations between H44 and platform elements H45 and uS11 in maps from the untreated particles (left). An example H45-/uS11-/H44+ volume is shown (right) and the red markers indicate the position of this volume in the scatterplots. (**C**) Results from performing principal component analysis on the voxels within a mask surrounding H44 for 500 volumes sampled from the H44+ subset of each dataset. The marginal distribution of the first principal component values from the two datasets is shown. (**D**) Volumes sampled along principal component 1 noted by colored markers in (C). (**E**) Molecular model of the untreated 30S_Δ*ksg*A_ structure with H44 in the inactive conformation. The inset highlights the linker helix that forms in this structure and contributes to stabilization the head and platform domains, with nucleotides important in stabilizing these domains annotated. (**F**) Molecular model of KsgA bound to the 30S_Δ*ksg*A_ particle depicting an equivalent region and in a similar orientation to that shown in the (E). Putative steric clashes that would exist between KsgA’s N-terminal domain and key rRNA helices are noted by semi-transparent rendering of these helices. (**G**) Integrated model depicting KsgA’s proposed role in late stage assembly of the ribosomal small subunit.

To better quantify the number of ribosomal particles in the inactive and active conformations we again employed our voxel PCA approach, now focusing on the H44 region. We observed that the first principal component cleanly segregated volumes on the basis of the inactive versus active conformation (**Figure 6C**), and that sampling volumes along this principal component allowed for visualization of the active-inactive transition (**Figure 6D, Supplementary Video 2**). By plotting the particle distribution along principal component 1 within each dataset we found that the inactive H44 conformation was over-represented in the untreated dataset, and that this inactive conformation effectively disappeared upon treatment with KsgA (**Figure 6C**). These observations were consistent with a role for KsgA in the pruning of the H44 inactive state and was reminiscent of classic proofreading systems (Hopfield, 1974).

### KsgA uncouples the head and platform domains by unfolding a key linker helix

To understand the mechanism by which KsgA prunes the inactive ribosomal particles and induces uncoupling of the head and platform domains, we built a molecular model (**Supplementary Table 1**) using the H44-inactive density map from the untreated dataset (**Supplementary Figure 3**; class 2). In our structure, residues 1397-1400 and 1502-1505 formed a small “linker helix” at the junction between body, platform, and neck (**Figure 6E**). We found that this linker helix was primed to position and stabilize helix 28 (H28), which is the main structural element determining the position of the head domain with respect to the body domain, and it also appeared to stabilize H24 and H45 in the platform domain (**Figure 6E**). In contrast, this linker helix and portions of H28 were absent in maps derived from KsgA-bound 30S_Δ*ksg*A_ particles, and the linker helix space was occupied by KsgA’s N-terminal domain (**Figure 6F**). This analysis suggested that upon binding, the N-terminal domain of KsgA disrupted this critical linker helix, thereby destabilizing the platform domain, inducing uncoupling of the head and body domains, and simultaneously displacing inactive conformations of H44.

Taken together, these data support a model in which 30S particles can assemble completely in the absence of KsgA, but in doing so produce a subset of inactive particles. Our results suggest that, when KsgA is present, it specifically targets these inactive particles, with KsgA binding resulting in partial subunit disassembly through undocking of inactive helix 44, destabilization of the platform domain, and uncoupling of the head and body domains upon destablization of a key linker helix in the subunit neck. We hypothesize that upon KsgA methylation and subsequent dissociation, these particles can then re-assemble with a new opportunity for helix 44 to adopt an active conformation, which, in totality should increase the overall fidelity of the assembly process (**Figure 6G**).

## DISCUSSION

### KsgA as an assembly factor

While KsgA’s non-essential but highly conserved role in rRNA methylation has been known for decades (Poldermans et al., 1979a; Poldermans et al., 1979b; Poldermans et al., 1979c; Sparling, 1970), recent genetic and biochemical assays have suggested an additional role for KsgA in supervising ribosome biogenesis (Connolly et al., 2008). Indeed, treatment of 30S subunit components with KsgA during *in vitro* reconstitution increases the activity of the resulting particles, but this effect is independent of the methylation activity (Cunningham et al., 1991; Cunningham et al., 1990; Igarashi et al., 1981). These results have led to a general model of KsgA acting as a late-stage ribosome biogenesis factor. In this model, KsgA is hypothesized to couple its binding to conformational rearrangements within the 30S that would allow that particle to more effectively undergo subunit joining and initiate translation (Connolly et al., 2008), with methylation serving to aid KsgA dissociation. Our study illuminates these long-hypothesized structural transitions, and reconciles decades of biochemical and genetic studies into an integrated model of KsgA’s role in late-stage ribosome biogenesis.

Specifically, our data reveal a “*proofreading*” role for KsgA, wherein it specifically recognizes subtly inactive subunits and, upon binding, displaces both the critical intersubunit helix 44 as well as an underappreciated linker helix that helps to stabilize the platform domain and to orient the head domain relative to the body. According to our structures, KsgA binding induces structural destabilization that uncouples the body, head, and platform domains, resulting in partial subunit disassembly. Our structures additionally confirm and highlight key atomic contacts that facilitate binding of substrate adenosines in the active site, and they provide a structural explanation for: 1) the order of base methylation; 2) how the rRNA contacts are maintained as successive substrates are flipped into the active site; and 3) how the methylated products are released. Taken together, these structures yield an assembly factor-mediated proofreading model in which nearly mature but inactive particles are recognized and destabilized by KsgA binding, resulting in partial subunit dissociation. We hypothesize that upon methylation and subsequent KsgA dissociation, these particles reassemble, and are thereby provided another opportunity to adopt a translationally active conformation (**Figure 6G**). We interpret this role of KsgA as a mechanism to maximize the efficiency of the assembly process for small ribosomal subunits, and to enforce the proper assembly order.

Finally, we note that our KsgA-bound structure, like others recently published (Schedlbauer et al., 2021; Stephan et al., 2021) is incompatible with subunit joining, explaining how expression of a catalytically inactive variant KsgAE66A profoundly inhibits cell growth (Connolly et al., 2008). Overall, this proposed role of KsgA resembles that recently assigned to RbgA, a ribosome assembly factor in Bacillus subtilis, which ensures the 50S subunit follows a canonical maturation pathway where the functional sites are the last structural motifs to mature (Seffouh et al., 2022).

### Uncovering coupled assembly reactions through structure

The classic Nomura assembly map (Held et al., 1974; Mizushima and Nomura, 1970), which depicts ribosomal protein binding inter-dependencies, has long guided our understanding of small subunit assembly. Incorporating the docking status of rRNA helices into such a map could, in principle, reveal new information linking assembly of ribosomal proteins and RNA helices. Indeed, generating such an extended Nomura map has long been a goal of the ribosome biogenesis field (Duss et al., 2019; Dutca and Culver, 2008; Jagannathan and Culver, 2003; Rodgers and Woodson, 2019; 2021; Stern et al., 1989; Woodson, 2011). However, experiments to determine such interdependencies are challenging, as one must assess the structural status of each rRNA helix with single-molecule resolution, and correlate its status with the presence or absence of each r-protein across the population. Nonetheless, we hypothesized that cryoDRGN’s powerful generative model, which we found can resolve rRNA helices and r-proteins in hundreds to thousands of density maps, might be well suited for such a task.

To test this hypothesis, we constructed a directed graph in which each node represented one rRNA helix or r-protein, and an edge between any two nodes reflected a dependency. To calculate inter-node dependencies, we used our subunit occupancy calculations from the KsgA-treated dataset and plotted the resulting dependency map (**Figure 7A**; Methods). Consistent with existing data supporting early formation of the ribosomal body relative to the head (Mulder et al., 2010), this analysis highlighted that primary binders – rRNA helices and r-proteins whose occupancy was independent of all other elements – were primarily located in the body (**Figure 2B**). In contrast, full occupancy of the head helices required both the body elements and rRNA helices 28, 35, and 36, which form the SSU “neck” (**Figure 7B; Supplementary Figure 12**). This analysis further highlighted the highly cooperative nature of head domain assembly. Specifically, sampled volumes exhibited either minimal or nearly complete occupancy for most head elements, resulting in no observable dependencies between head r-proteins uS3, uS14, uS9, and uS7, and the core head helices (H29, H30, H31, H37-H43). In contrast, the rRNA helices at the most extreme terminus of the head (H32 and H33) appeared to depend on the formation of the core head elements, and indirectly on helix 28. We interpret this dependence as reflecting the mobility of the undocked head in many KsgA-treated particles; this interpretation is consistent with the voxel PCA analysis that showed an increase head mobility upon KsgA treatment. Taken together, this novel analysis allowed for an expansion of the classic ribosome assembly maps to now include rRNA helices, and we expect that such maps can be further refined as additional assembly intermediates are structurally characterized.

**Figure 7:**
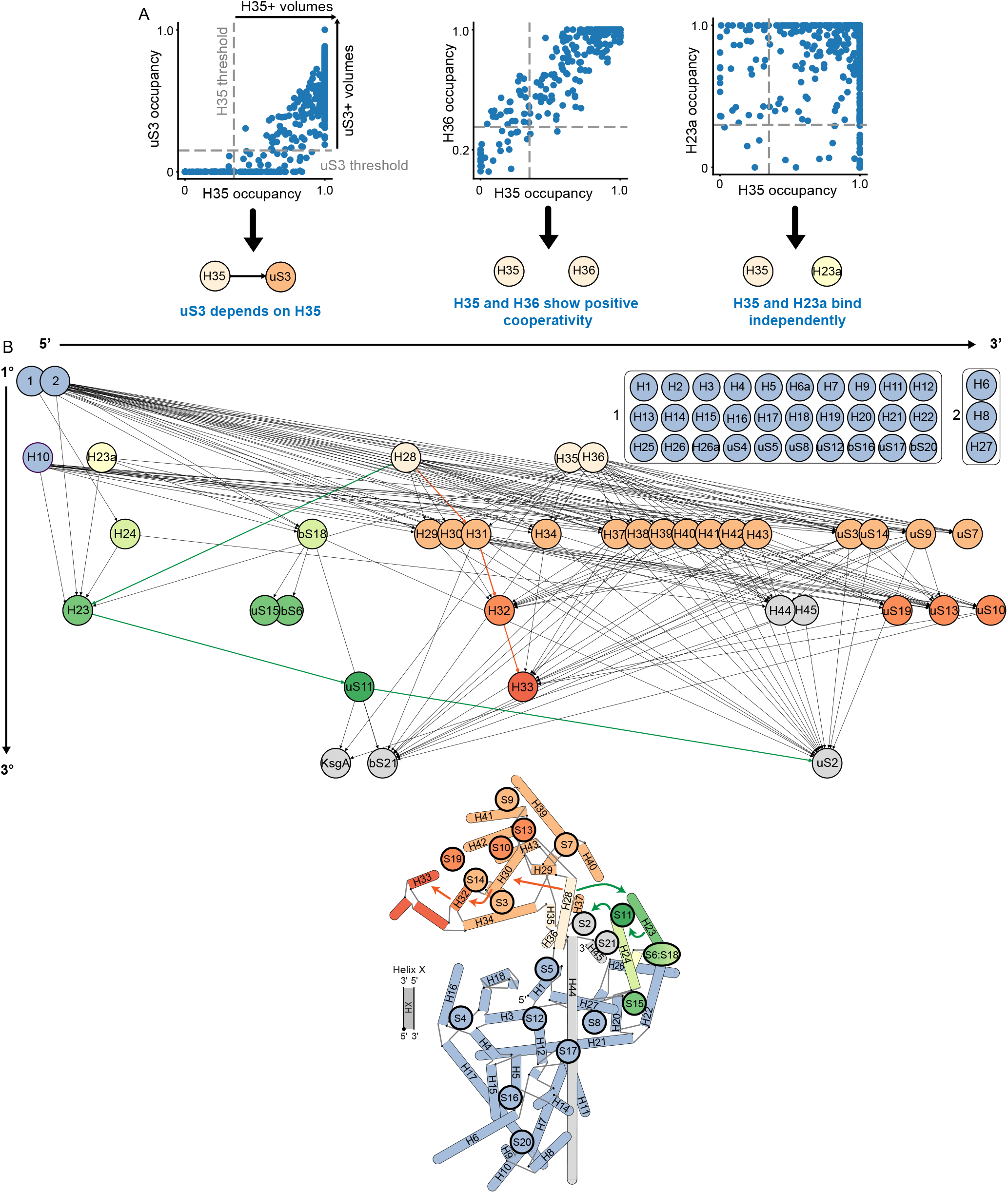
Network analysis reveals assembly dependency map for KsgA-treated SSUs. (**A**) Examples of dependency relationship calculations, as described in Methods. Dependency relationships were defined as a unidirectional requirement of occupancy of a given subunit for occupancy of another. (**B**) A directed acyclic graph constructed from the calculated dependency relationships, in which each node is an r-protein or rRNA helix, and each dependency relationship is a directed edge from the independent subunit to the dependent subunit. Edges of the graph were pruned to eliminate direct paths between any two nodes if there existed also an indirect path between these nodes. Nodes were also consolidated (boxed nodes) if they had all the same incoming and outgoing edges. With the exception of the consolidated body elements, nodes are arrayed horizontally 5’-to-3’ along the rRNA transcript and positioned vertically to reflect primary, secondary, and tertiary elements as determined by this graphical analysis. Nodes are colored by domain in line with the model below, which was modified from Sykes & Williamson 2009. Colored arrows highlight representative key links between helix 28 and downstream head and platform domain elements.

### Systematic characterization of highly heterogeneous structural ensembles

Traditional approaches to resolve structural heterogeneity employ iterative rounds of hierarchical 3D classification, and require significant expert-guided intervention, with users supplying the number of classes within each round, as well as the total number of rounds of classification. Although recent work has proposed quantitative standards for determining when a dataset has been sufficiently classified (Rabuck-Gibbons et al., 2022), questions remain about how to choose the number of classes or the classification end-point, and how robust results are to these classification parameters. It is thus desirable to develop methods to analyze and quantify structural heterogeneity that are more unbiased, reproducible, and quantitatively robust. The cryoDRGN training and systematic subunit occupancy analysis approach we present here represents one avenue for conducting this type of quantitative analysis.

Comparing the results of traditional 3D classification and subunit occupancy analysis on the datasets presented here suggests that grossly similar classes of particles are identified by the two approaches (**Figure 1, Supplementary Figure 13**). However, the subunit occupancy analysis of cryoDRGN generated maps was able to identify and quantify heterogeneity on a more granular scale than permitted by 3D classification, allowing maps that differed only by the presence or absence of single proteins to be identified (**Figure 2A-B**). Notably, guided by this cryoDRGN analysis, one can readily identify particle subsets that, when analyzed with traditional tools, produce reconstructions mimicking those from cryoDRGN (**Supplementary Figure 13**, see Methods). The subunit occupancy analysis also identified rare structural states missed by traditional classification, including a small subpopulation of immature KsgA-bound 30S particles completely lacking density for the head (**Figure 3B**). Furthermore, by sampling hundreds of volumes from the structural ensemble, the subunit occupancy analysis has greater statistical power to extract correlative relationships between any two given subunits, which we used to identify individual binding prerequisites for KsgA (**Figure 3D**).

Occupancy analysis nonetheless has several key limitations – principally that it is an atomic model-based approach and relies on the assumption that subunits are either present in their native conformation, or absent. As exemplified by helix 44 and the head domain, this approach is challenged by conformational heterogeneity, where subunits may be present but in an alternative location. In such instances, we found applying a voxel-based principal component analysis to relevant subsets of the particle stack was a powerful approach to characterize the local motions of conformationally flexible subunits (**Figure 6C**).

In addition to elucidating the role of KsgA in ribosome biogenesis, the approaches outlined here for systematically analyzing and quantifying structural landscapes may prove more broadly useful in realizing the single-molecule potential of cryo-EM. In combination with the various machine learning-based approaches recently developed for generating large volume ensembles from cryo-EM datasets (Chen and Ludtke, 2021; Zhong et al., 2021), our subunit occupancy analysis and voxel-based principal component analysis approaches provide new tools to leverage the heterogeneity present in single particle cryo-EM datasets to uncover biological insights about dynamic proteins and protein complexes.

## MATERIAL AND METHODS

### Bacterial strains and protein overexpression clones

The parental *Escherichia coli* K-12 (BW25113) and *ksg*A null (JW0050-3) strains from the Keio collection (Baba et al., 2006) were obtained from *E. coli* Genetic Resource Center, Yale University. The sequence of the *ksg*A gene (NCBI reference sequence: NC_000913.3) with a thrombin-cleavable N-terminal His_6_ tag was optimized for over expression in *E. coli* cells using GeneOptimizer software, synthesized (Life Technologies; Thermo Fisher Scientific), cloned into the carrier pMA-T plasmid using the SfiI and SfiI cloning sites, and subsequently subcloned into the final expression vector pET15b using the NdeI and a BamHI restriction sites.

### Purification of ribosomal particles

The immature 30S_Δ*ksg*A_ subunits were purified from *E. coli ksg*A Keio collection (Baba et al., 2006) deletion strain JW0050-3; 30S_WT_ subunits were purified from parental strain BW25113. All strains were grown in LB (4L for JW0050-3; 3L for BW25113) at 25°C with shaking (225 rpm). Cells were cooled to 4°C and collected by centrifugation at 3,000g for 15 mins in a Beckman JLA-8.1000 rotor upon reaching OD600=0.2 (JW0050-3) or OD600=0.6 (BW25113). Pellets were then resuspended in 14 mL buffer A [20 mM Tris-HCl at pH 7.5, 10 mM magnesium acetate, 60 mM NH_4_Cl, 0.5 mM EDTA, 3 mM 2-mercaptoethanol, cOmplete protease inhibitor tablet (Roche), DNase I (Roche)]. Resuspended cells were lysed by sonication on ice and the cell lysate was centrifuged at 42,000g for 30 minutes in a Beckman 70Ti rotor to clear cell debris. The supernatant was layered over a 1.1 M sucrose cushion in buffer A lacking protease inhibitors (3 mL supernatant and 3mL sucrose cushion) and centrifuged for 16 hours at 118,000g in a Beckman 70Ti rotor. The pellet containing the ribosomal particles was then resuspended in buffer E [10 mM Tris-HCl at pH 7.5, 10 mM Mg acetate, 60 mM NH_4_Cl, 3 mM 2-mercaptoethanol] (30S_Δ*ksg*A_) or buffer C [10 mM Tris-HCl pH 7.5, 10 mM Mg acetate, 500 mM NH_4_Cl, 0.5 mM EDTA, and 3 mM 2-mercaptoethanol] (30S_WT_). Resuspended crude ribosomes (∼120 A260 units) were applied to 34 mL 10%–30% (w/v) sucrose gradients prepared in buffer E. Gradients were then centrifuged at 31,000g for 16 hours in a Beckman SW32Ti rotor and fractionated using a Brandel fractionator apparatus and an AKTA Prime FPLC system (GE Healthcare). The profile was monitored by UV absorbance at A254 and the relevant fractions were collected. Fractions for each ribosomal particle were pooled and spun down in a Beckman MLA-80 rotor for 16 hours at 108,000g. The resulting pellets (30S_Δ*ksg*A_) were washed and resuspended in 75 μL buffer E, flash frozen in liquid nitrogen, and stored at -80°C. Washed 70S_WT_ pellets were resuspended in buffer F [10 mM Tris-HCl, pH 7.5, 1.1 mM magnesium acetate, 60 mM NH_4_Cl, 0.5 mM EDTA, and 2 mM 2-mercaptoethanol], and ∼120 A260 units were applied to a 34 mL of 10%–30% (wt/vol) sucrose gradient prepared with buffer F. The gradient was centrifuged and fractionated, as above. Fractions containing 30S_WT_, which resulted from 70S_WT_ dissociation in buffer F, were selected, pooled and pelleted as above. They were then resuspended in 200 μL buffer E, flash frozen, and stored at -80°C. Mature 70S ribosomes used for analysis by mass spectrometry were purified from the parental strain BW25113 as previously described (Razi et al., 2019).

### Protein overexpression and purification

KsgA was purified from *E. coli* strain BL21-A1 transformed with the pET15b-ksgA plasmid. Cells were grown at 37°C with shaking (225 rpm) in LB medium supplemented with 100 μg/mL ampicillin; expression was induced at an OD600 of 0.6 by adding L-arabinose (0.2%) and IPTG (1mM) and incubating at 25°C for 4 hours. Cells were harvested by centrifugation at 3,700g for 15 minutes and washed with 30 mL of PBS 1x buffer before resuspension in 20 mL of buffer A1 [50 mM Na_2_HPO_4_ at pH 7.5, 300 mM NaCl, 5% glycerol], with addition 1 mM PMSF, 1 mM benzamidine, 5 μg/ml leupeptin, 70 μg/ml pepstatin. The cells were lysed by sonication on ice, then centrifuged at 30,000g for 45 minutes to clear cell debris. The supernatant was filtered with a 0.45-μm syringe filter (Millipore) and loaded onto a HisTrap HP column (GE Healthcare). The column was washed with eight column volumes of buffer A1 containing 75 mM imidazole and six column volumes of buffer A1 containing 100 mM imidazole. KsgA was eluted with 250 mM imidazole in buffer A1. Purity of the fractions was monitored by SDS/PAGE and fractions containing KsgA protein were collected and pooled together. The N-terminal His_6_ tag was removed by digestion with thrombin (GE Healthcare) by adding the enzyme at a concentration of 10 Units per mg of KsgA protein during overnight dialysis against PBS. Precipitated protein was removed by filtration, and the filtrate was loaded on to a Hi Trap SP HP column (GE healthcare) equilibrated in buffer B1 [50 mM Na_2_HPO_4_ at pH 7.5, 50 mM NaCl, 5% glycerol]. The column was washed with ten column volumes of buffer B1 containing 80 mM NaCl and then eluted with buffer B1 containing 340 mM NaCl. Fractions containing KsgA were pooled and concentrated using a 10-kDa cutoff filter (Amicon), and the concentrated KsgA was then diluted in storage buffer [50 mM Na_2_HPO_4_ at pH 7.5, 50 mM NaCl, 5% glycerol] at the ratio of 1:10 before storage at -80°C.

### Microscale thermophoresis experiments

The amine residues of purified KsgA were fluorescently labeled with NHS red using the Protein Labeling Kit RED-NHS 2nd Generation (Cat # MO-L011 Nanotemper). The labeling reaction was performed according to the manufacturer’s protocol by mixing KsgA at a final concentration of 20 μM with a 3-fold molar excess of dye at room temperature for 30 minutes in the dark. The provided labeling buffer was supplemented with 10 mM magnesium acetate. Free dye was eliminated using the Gravity Flow Column B pre-equilibrated with buffer containing 10 mM Tris-HCl pH 7.5, 15 mM MgCl_2_, 6 mM 2-mercaptoethanol and 0.05% Tween 20. Labeled KsgA was diluted in MST buffer [10mM Tris-HCl pH 7.5, 60mM NH_4_Cl, 15 mM Mg acetate, 6 mM 2-mercaptoethanol, 0.05% Tween 20] to a concentration of 100 nM, and a serial dilution of ribosomal particles in MST buffer was prepared. The labeled KsgA was mixed 1:1 (v/v) with each different concentration of ribosomal particles, yielding a final concentration of KsgA at 50 nM, and concentrations of ribosomal particles spanning from 0.053 nM to 1.75 μM. All reactions were incubated for 20 minutes at 25 ºC before loading into premium glass capillaries (NanoTemper Technologies). Microscale thermophoresis (MST) measurements were performed using the Monolith NT.115 microscale thermophoresis instrument (NanoTemper Technologies) at 25 ºC. Experiments were conducted at LED power of 100% and medium MST IR-laser power. The resulting binding curves and dissociation constants (KD) were obtained by plotting the normalized fluorescence (Fnorm = F_1_ / F_0_) versus the logarithm of the ribosomal subunit concentration. The obtained K_D_ values were calculated from three independently performed experiments using the NanoTemper analysis software (version 2.2.6).

### Quantitative mass spectrometry analysis

For mass spectrometry analysis of KsgA-treated immature 30S_Δ*ksg*A_ particles, KsgA (10 μM) was mixed with 30S_Δ*ksg*A_ particles (1 μM) in a 200 μL reaction in modified buffer E containing 6 mM 2-mercaptoethanol, and the reaction mixture was incubated at 37°C for 20 minutes. 50 μL aliquots of each reaction were laid over a 150 μL 1.1 M sucrose cushion in buffer E and then ultra-centrifuged at 436,000g for 3.5 hours in a Beckman Coulter TLA-100 rotor. The pellets were resuspended in 20 μL of buffer E and concentration was measured at A260 prior to flash freezing in liquid nitrogen and stored at -80°C. The sample containing untreated 30S_Δ*ksg*A_ particles was prepared in a similar manner, but KsgA was not added to the initial reaction.

Samples were prepared in triplicate for mass spectrometry by resuspending 10 pmol of each sample in ribosome lysis buffer [20 mM Tris, pH 7.6, 200 mM NH_4_Cl, 0.5 mM EDTA, 10 mM MgCl_2_, 6 mM 2-mercaptoethanol, 13% trichloroacetic acid] and spiking each resulting sample with a constant volume of ^15^N-labeled cellular lysate that was previously assessed to provide roughly stoichiometric quantities of ribosomal proteins for normalization. Samples were incubated on ice for 30 minutes, then centrifuged for 30 minutes at 4°C and washed with 10% TCA and acetone. Following the final wash, pellets were dried at room temperature for 30 minutes, then resuspended in 100 mM Na_4_HCO_3_ and 5% acetonitrile. Samples were reduced by adding dithitreitol (5 mM) and incubating in a 65°C water bath for 10 minutes, then alkylated by adding iodoacetamide (10 mM) and incubating at 30°C for 30 minutes. Trypsin digestion was carried out overnight at 37°C, then samples were desalted using Pierce C18 spin columns.

For each sample, peptides were spiked with Pierce iRT standards (450 fmol) and then were loaded in buffer MSA [4% acetonitrile, 0.1% formic acid] onto an Acclaim PepMap 20mm C18 column coupled to an EASYSpray nano 500mm analytical column (Thermo) through a switching valve. After washing with MSA, peptides were eluted from the analytical column across a 90 minute 4%-40% gradient of acetonitrile in MSA and injected onto a Q-Exactive HF-X mass spectrometer (Thermo). Data was collected in replicate either in variable-window DIA or top-12 DDA acquisition modes. DIA acquisitions used the following parameters: 70 variably spaced MS^2^ isolation windows spanning 390-1390 Thompsons with 25 NCE collision energy, 35 ms max injection time, 5e5 AGC target, and 15k resolution, with 3 MS^1^ scans over the range 390-1390 Thompsons collected at 120k resolution, 3e6 AGC target, 35 ms max injection time evenly interspersed over the 70 MS2 scans each cycle. Top-12 DDA acquisitions used the following parameters: MS^1^ acquisition at 60k resolution, 3e6 AGC target, 50 ms max injection time scanning 390-1390, and MS^2^ acquisitions at 15k resolution, 25 NCE collision energy, 1e5 AGC target, 100 ms max injection times, and 2 Thompson isolation windows.

DDA results were pooled and searched with Comet and iProphet to create a library for searching DIA data. DIA data was manually curated to select high signal peaks in Skyline, and the resulting report exported for normalization. Peptide abundances were normalized to the intensity of the ^15^N peak, and protein intensity calculated as the median normalized MS^1^ peptide intensity. The stoichiometry relative to the wildtype 70S ribosome was calculated for each protein by dividing the protein intensity in the given sample by the median protein intensity in the wildtype 70S samples. The results of these stoichiometry calculations were then hierarchically clustered.

### Cryo-electron microscopy

Immature 30S_Δ*ksg*A_ particles were diluted in modified buffer E that contained 6 mM 2-mercaptoethanol to a final concentration of 720 nM. For KsgA-treated immature 30S_Δ*ksg*A_ particles, KsgA was added in a 10-fold excess to obtain a solution with ribosomal subunits and KsgA at concentrations of 0.5 μM and 5.2 μM, respectively. Both samples were incubated at 37 ºC for 20 minutes before sample vitrification was performed in a Vitrobot Mark IV (Thermo Fisher Scientific Inc.) at 25 ºC and 100% humidity. For all grids, 3.6 μL of the relevant sample was applied to holey carbon grids (C-flat CF-2/1–3Cu-T) that had been glow discharged in air at 15 mA for 15 seconds. Grids were blotted for 3 seconds with a blot force of +1 before plunging.

Datasets for the immature 30S_Δ*ksg*A_ subunits and KsgA-treated immature 30S_Δ*ksg*A_ were collected using SerialEM software (Schorb et al., 2019) in the Titan Krios at FEMR-McGill (**Supplementary Table 1**). Movies were recorded in a Gatan K3 direct electron detector equipped with a Quantum LS imaging filter. The total dose used for each movie was 45 e/Å2 equally spread in 33 frames for the untreated dataset, and 71 e/Å2 total dose across 30 frames for the KsgA-treated dataset. Both datasets were collected at a magnification of 105,000x, yielding images with a calibrated pixel size of 0.855 Å. The nominal defocus range used during data collection was between -1.25 and -2.75 μm.

### Image processing with RELION

Cryo-EM movies were corrected for beam induced motion using RELION’s implementation of the MotionCor2 algorithm (Zheng et al., 2017; Zivanov et al., 2018). CTF parameter estimation was done using the CTFFIND-4.1 program (Rohou and Grigorieff, 2015) and the remaining processing steps were done using RELION 3.1.2 (Zivanov et al., 2018). In the untreated dataset, a total of 775,859 particles were selected with auto-picking and subsequently extracted. This particle stack was subjected to two cycles of reference-free 2D classification to remove particles incorrectly selected by the auto-picking step. These steps resulted in a dataset comprised of 552,604 particles. To separate the particles representing the various assembly intermediates of the 30S_Δ*ksg*A_ particles, we performed a multi-layered 3D classification strategy that resulted in the seven classes shown in **Supplementary Figure 3A**. The initial 3D reference used for these classifications was obtained by the random sample consensus (RANSAC) approach as implemented in Scipion (Gomez-Blanco et al., 2019). Resulting maps from 3D classification steps were visually inspected in Chimera (Pettersen et al., 2004) and those particles assigned to classes representing the same assembly intermediate were pooled together for high-resolution refinement.

All maps for the various classes were refined in four steps: In the first step, the 3D auto-refine process was performed without a mask, using as an initial model the maps obtained via 3D classification after a 60 Å low-pass Fourier filter was applied. The resulting maps were used to create a mask and used as the initial model for a second step of refinement. The outputs of the last 3D auto-refine process were then used as input for CTF refinement, using default parameters with the additional estimation of beam tilt. Finally, Bayesian polishing was performed to correct for per-particle beam-induced motion before subjecting these particles to a final round of 3D refinement. Bayesian polishing was performed using sigma values of 0.2, 5,000 and 2 for velocity, divergence and acceleration, respectively. Sharpening of the final cryo-EM maps was done with RELION. Resolution estimation is reported using an FSC threshold value of 0.143. Local resolution analysis was done with RELION. Cryo-EM map visualization was performed in UCSF Chimera (Pettersen et al., 2004) and Chimera X (Goddard et al., 2018; Pettersen et al., 2021).

The smaller KsgA-treated dataset, which generated the cryo-EM maps of the various classes shown in **Supplementary Figure 3B**, and the larger dataset that yielded the high-resolution cryo-EM structure of the KsgA-bound 30S_Δ*ksg*A_ complex in **Figure 4**, were both processed using the same pipeline as described above. The initial number of particles extracted after auto-picking were 588,015 and 1,821,260 particles, respectively, in these datasets. After two cycles of reference-free 2D classification for each dataset, we produced two particle stacks containing 369,621 and 665,547 particles that were processed separately. To best define the density representing KsgA in the class exhibiting KsgA bound, particles in this group were subjected to a focused 3D classification using a spherical mask around the KsgA binding region. Particles in the class with clearer KsgA density were subjected to one additional cycle of 3D auto-refine, resulting in 231,280 particles contributing to the high-resolution cryo-EM map of the KsgA-bound 30S_Δ*ksg*A_ complex shown in **Figure 4** and **Supplementary Table 1**.

### CryoDRGN training

Neural network analysis of structural heterogeneity was carried out using cryoDRGN v0.2.1 and v0.3.2b (Zhong et al., 2021). For both KsgA-treated and untreated datasets, the full particle stacks from RELION AutoPicking were run through ab initio model generation and 3D refinement with cryoSPARC (Punjani et al., 2017). These consensus reconstructions were used to supply the poses for an initial round of low-resolution cryoDRGN training, in which particles were downsampled to a box size of 128 (2.5367 Å per pixel). The networks for both datasets were trained with a 10-dimensional latent variable and 1024×3 encoder and decoder architectures.

After 50 epochs of low-resolution training for the KsgA-treated dataset and 46 epochs of training for the untreated dataset, the particle stacks were filtered to exclude 70S ribosomes, edge artifacts, ice contaminants, and particles that led to poor quality 3D reconstructions. For the KsgA-treated dataset, filtration was implemented by selecting particles that satisfied the following criteria after UMAP (Becht et al., 2018) dimensionality reduction: UMAP2 < (−2.5*UMAP1 + 15). Filtering reduced the size of the particle stack from 588,015 to 267,905 particles. Likewise, the untreated dataset, consisting of 775,859 particles, was filtered by selecting particles with UMAP2 < (UMAP1 - 0.8), resulting in a final stack of 394,110 particles. These filtered particle stacks were returned to Relion for joint refinement and Bayesian polishing as described above.

To improve pose assignments, the particles were extracted after Bayesian polishing and the particle stack was imported into cryoSPARC for ab initio model generation and 3D refinement. Model generation and 3D refinement were done both with all particles combined, and with each of the KsgA-treated and untreated dataset particles individually. Pose assignments extracted from the cryoSPARC refinements were paired with the RELION CTF parameters and particle stacks for an additional round of cryoDRGN training in which particles with poor pose assignments were filtered. This filtration training was done at box size 256 (1.2684 Å per pixel), with an eight-dimensional latent variable, and a 1024×3 architecture for both the encoder and decoder networks.

Datasets trained individually were filtered by the magnitude of the latent variable, whereas the co-trained dataset was filtered by eliminating particles within k-means clusters visually determined to represent poor pose assignments. Filtering reduced the size of the KsgA-treated dataset to 250,325 particles (retaining 93.4% of the particles) and reduced the size of the untreated dataset to 364,289 particles (retaining 92.4% of the particles). Filtering of the co-trained dataset eliminated 65,168 particles of 662,015 total (retaining 98.4% of the particles). These final filtered particle stacks were subjected to a final round of high resolution cryoDRGN training, with a box size of 256 and eight-dimensional latent variable, and 1024×3 architecture for both the encoder and decoder networks. The full cryoDRGN filtration and analysis pipeline is presented in **Supplementary Figure 6A**.

### Subunit occupancy analysis

For subunit occupancy analysis, 500 volumes were systematically sampled at k-means cluster centers of the latent embeddings. Existing atomic models of the ribosome – PDB: 4V9D, (Dunkle et al., 2011); PDB: 4ADV, (Boehringer et al., 2012) – were used to create masks corresponding to each of the rRNA helices and ribosomal proteins, as well as KsgA. Each of these 69 masks was applied to each of the 500 volumes in turn, and the intensities of all voxels within each masked region was summed. The summed voxel intensity measurements were normalized by the summed voxel intensities of the corresponding subunit found in a map generated from the atomic model using the molmap tool in Chimera (Pettersen et al., 2004), producing a fractional occupancy measurement. Fractional occupancies were then scaled from the 10^th^ to the 70^th^ percentile of the dataset, and hierarchically clustered to identify patterns in subunit occupancy. Volume classes and structural blocks were defined by setting a threshold distance in the hierarchical clustering. Centroid volumes for each volume class were generated by calculating the median z-coordinates of all particles in the relevant class and identifying the nearest neighbor particle in the original stack to this median point. The code used to carry out these analyses is available at https://github.com/lkinman/occupancy-analysis.

Volumes from the original hierarchical clustering with a fractional KsgA occupancy of greater than 0.15 were designated as KsgA-bound and were isolated for another round of hierarchical clustering to produce the KsgA-bound heatmap. Similarly, volumes from the original clustering were designated as H44+ if the fractional occupancy of H44 was greater than 0.1. All particles in the corresponding k-means clusters were additionally designated as H44+, and 500 new volumes were generated from k-means centroid locations of latent space defined by the H44+ subset of each of the datasets. PCA-based analysis of these volumes was done by amplitude-scaling all the volumes relative to a representative volume from the lower-resolution KsgA dataset with Diffmap (http://grigoriefflab.janelia.org/diffmap), aligning the untreated volumes to the KsgA-treated volumes with EMAN2 (Tang et al., 2007), applying a 12 Å-extended mask around the H44 region, and finally performing PCA on the resulting voxel array. A similar approach was taken for voxel analysis of the head domain, designating all volumes in classes 1-6 in the KsgA-treated dataset, and all classes except class 8 in the untreated dataset, as head+, and sampling 500 new volumes from the head+ subset of latent space. Volumes were again amplitude-scaled to a common map from the KsgA-treated dataset and aligned. A mask was applied to the H32/H33 region, and PCA performed on the resulting voxel array.

To calculate the summed squared residual (SSR) between each sampled volume and a mature 30S volume, we first downsampled each of the 500 samples maps from each dataset to a boxsize of 64. The volumes were then binarized, and the only voxels used for the SSR calculations were those occupied in at least 1% of the relevant volume ensemble. The SSR was calculated over this subset of voxels between each sampled volume and the relevant mature centroid volume (class 3 in each dataset).

### Nomura assembly map analysis

Determination of occupancy dependency relationships between subunits was done by defining a threshold for each subunit that divides low-occupancy volumes from high-occupancy volumes. The thresholds were set based on expert-guided manual inspection of the volumes above and below the threshold. For any given subunit s and associated occupancy threshold ts, we define H(s) to be the set of volumes with occupancy of s greater than ts. We calculate the fractional dependency of s on any other subunit r as:

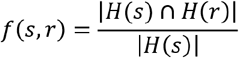

A directed edge *e*_*rs*_ from *r* to *s* is built if *f(s,r)*≥0.85 and *f(r,s)*<0.9. The resulting directed acyclic graph is then pruned by eliminating each edge *e*_*rs*_ if there exists another path from *r* to *s*. Finally, all nodes with identical in- and out-edges were grouped and treated as a single node in the resulting graph, as these nodes cannot be distinguished by our graphical analysis.

### Cryo-EM map analysis and molecular model building

To build the molecular model of the KsgA + 30S_Δ*ksg*A_ complex, we first performed multi-body refinement by dividing the consensus cryo-EM map into three major bodies (body 1: 30S body, body 2: 30S platform + KsgA and body 3: 30S head). Then, a soft mask was generated for each body and applied to the corresponding map during refinement. We used the automatic sharpening tool’phenix. auto sharpen’ from Phenix (Adams et al., 2010) to further improve the connectivity of the three cryo-EM maps derived from the multi-body refinement process. The molecular model was built from the available structures of the mature 30S ribosomal subunit (PDB: 4YBB) (Noeske et al., 2015) and *E. coli* KsgA (PDB: 1QYR) (O’Farrell et al., 2004). The atomic model of the 30S subunit was truncated into three domains that matched the three bodies in the multi-body refinement process. These initial models were fit into the cryo-EM maps by rigid-body docking in Chimera (Pettersen et al., 2004). The model for each body was built independently by successive rounds of real space refinement in Phenix (Adams et al., 2010) and manual model building in Coot (Emsley and Cowtan, 2004; Emsley et al., 2010). The three resulting molecular models were docked into the consensus KsgA-treated 30S_Δ*ksg*A_ complex cryo-EM map using the ‘dock_in_map’ tool in Phenix (Adams et al., 2010). Amino acids in the protein components and nucleotides in the 16S rRNA at the interfaces between the 3 bodies were manually built based on the density in the consensus map using Coot. Finally, the molecular coordinates from the three bodies were combined into the entire model for the KsgA-treated 30S_Δ*ksg*A_ complex using Coot (Emsley and Cowtan, 2004; Emsley et al., 2010). The cryo-EM maps for the three bodies of the KsgA-treated 30S_Δ*ksg*A_ complex obtained from multi-body refinement were rigid body fit to the consensus map and combined into a single high-resolution consensus map using ‘vop add’ command in Chimera.

We also performed multi-body refinement of the cryo-EM map before building a model for the untreated 30S_Δ*ksg*A_ particle with the helix 44 in the inactive conformation. In this case, the consensus map was divided into two bodies (body 1: body and platform and body 2: head). The molecular model for this structure was built using the same approach as that obtained for the KsgA-treated 30S_Δ*ksg*A_ complex. However, in this case the molecular model was built using PDB model 7BOF (Schedlbauer et al., 2021) as starting point.

## DATA AVAILABILITY

The density map and the model for the KsgA-bound 30S_Δ*ksg*A_ and untreated 30S_Δ*ksg*A_ structures will be deposited in the Electron Microscopy Data Bank (EMDB) and in the Protein Data Bank (PDB) upon publication, with unfiltered particle stacks deposited at EMPIAR. Trained cryoDRGN models will be deposited at Zenodo. The described occupancy analysis tools are available at: https://github.com/lkinman/occupancy-analysis.

## ACKNOWLEDGEMENTS

We thank K. Sears, M. Strauss, K. Basu and other staff members of the Facility for Electron Microscopy Research (FEMR) at McGill University for help with microscope operation and data collection; MIT-Satori administrative team for providing computational resources and support; and B. Powell and E. Zhong, and other members of the Davis and Ortega labs for constructive feedback on this work.

---

**Supplementary Figure 1.**
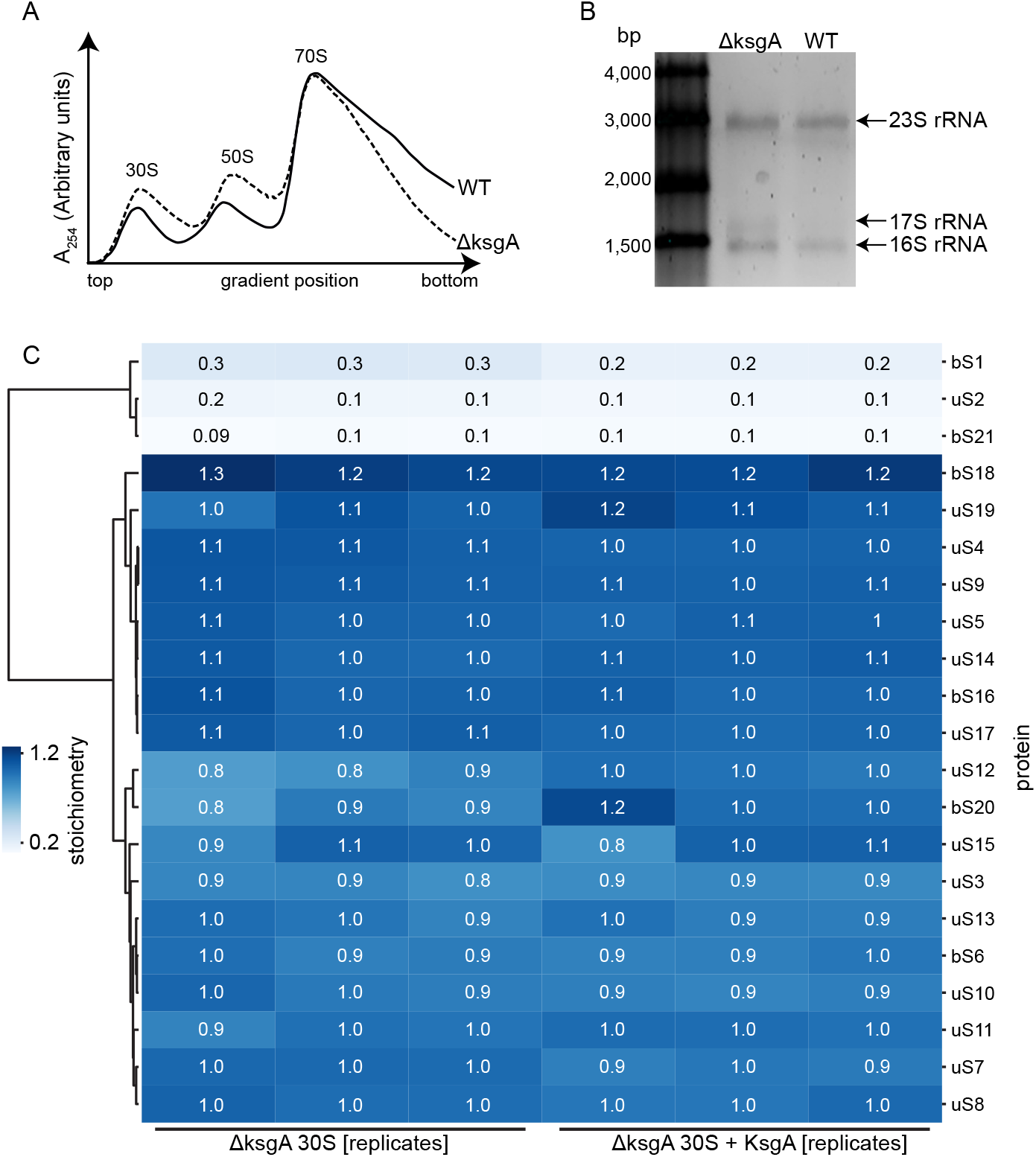
Purification and compositional analysis of 30S_Δ*ksg*A_ particles. (**A**) Sucrose gradient profiles from wild-type and Δ*ksg*A E. coli cells grown at 25 ºC. Peaks containing the 30S, 50S and 70S particles, and fractions used for this study are indicated. (**B**) Total rRNA from the wild type and the Δ*ksg*A cells purified and resolved by agarose gel electrophoresis, with relevant rRNA species noted. (**C**) Clustered heat map of protein abundance of KsgA-treated and untreated 30S_Δ*ksg*A_ particles determined by quantitative mass spectrometry (see Methods). In each replicate, protein abundance is reported as the stoichiometry relative to that measured in the wildtype 70S ribosome.

**Supplementary Figure 2.**
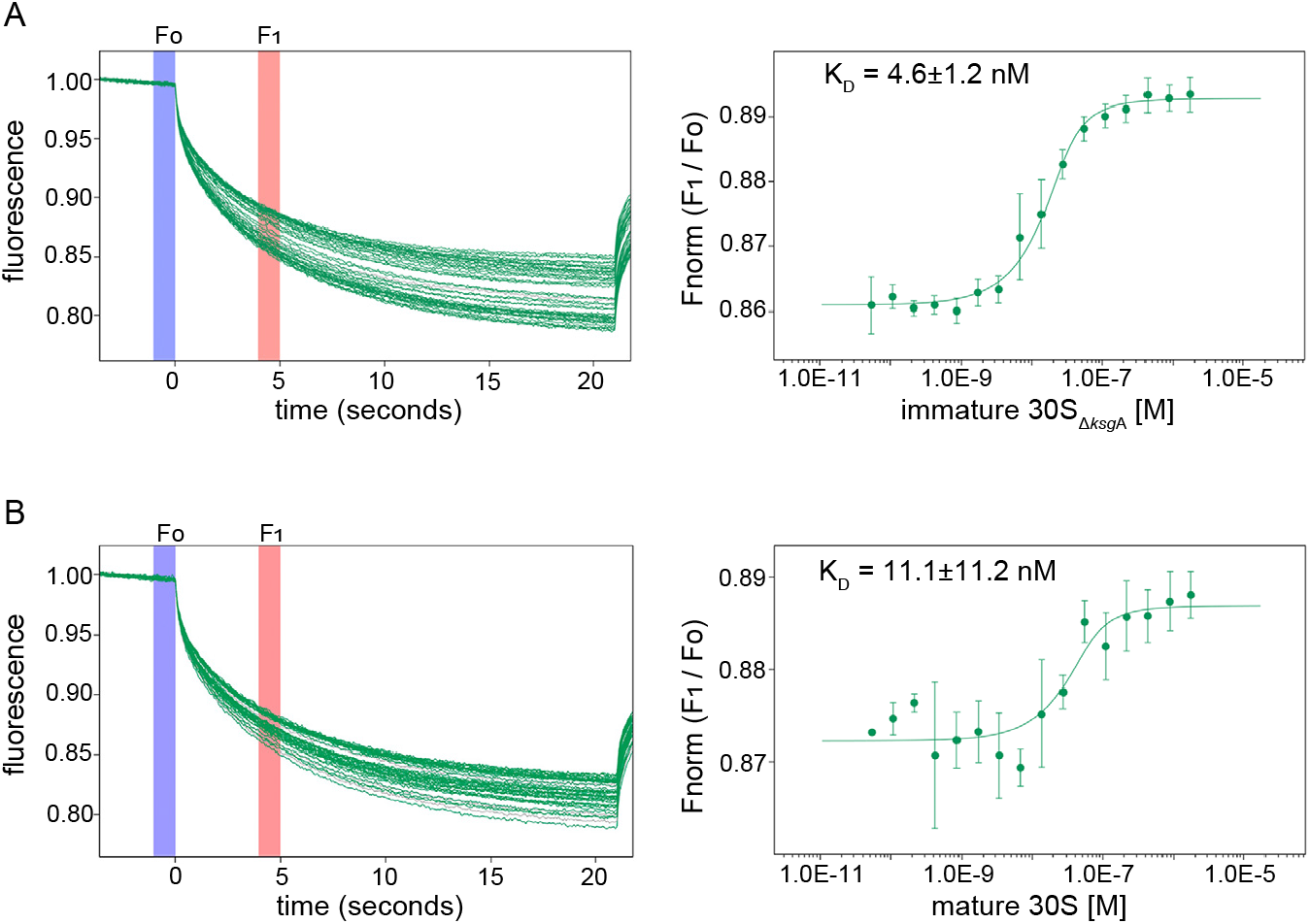
Binding affinity of KsgA to ribosomal particles measured by microscale thermophoresis (MST). Each reaction contained 50 nM fluorescently labeled KsgA and increasing concentrations of the immature 30S_Δ*ksg*A_ particles **(A)** or mature 30S particles that were derived from dissociated 70S_WT_ particles **(B)**. Thermophoretic mobility traces of the MST reactions (left) depict individual traces for each ribosomal particle concentration, and highlight the F_0_ (blue) and F_1_ (red) regions used to calculate binding. KsgA binding plots (right) depict Fnorm (F_1_/F_0_) versus particle concentration. The Fnorm curves were fit using the law of mass action to derive K_D_ values, which are reported with a 68% confidence interval derived from the variance of the fitted parameter. Error bars mark standard deviation from three replicate measurements at each concentration.

**Supplementary Figure 3.**
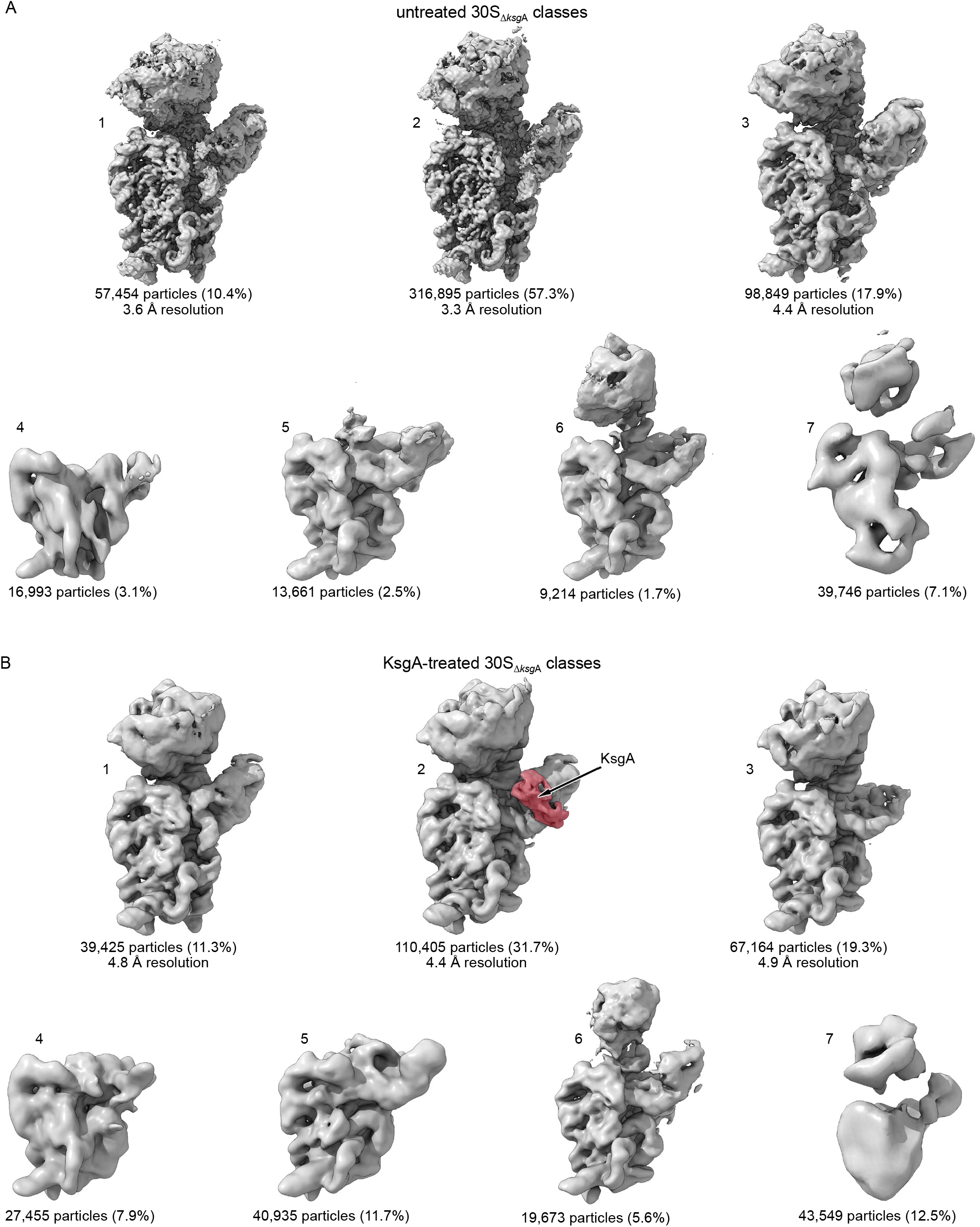
Discrete classification of the immature 30S_Δ*ksg*A_ particles before and after treatment with KsgA. 3D density maps of **(A)** untreated or **(B)** KsgA-treated 30S_Δ*ksg*A_ particles resulting from traditional 3D classification and reconstruction in RELION (Zivanov et al., 2018). Particle count and percentage of original dataset assigned to each class is noted. When high-resolution maps were obtained gold-standard FSC resolution estimate is noted. All maps are unsharpened.

**Supplementary Figure 4.**
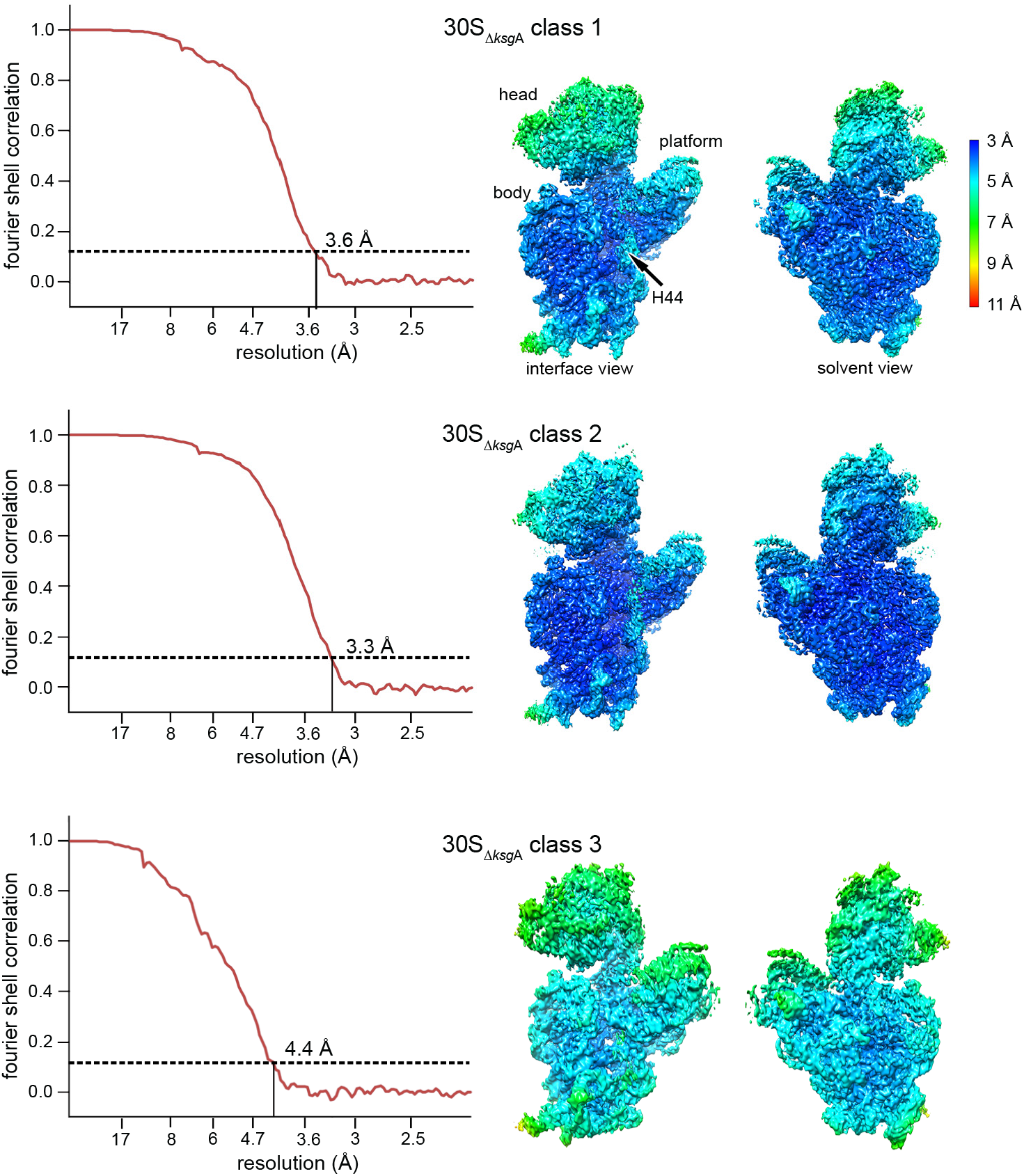
Resolution analysis of untreated 30S_Δ*ksg*A_ cryo-EM maps. Fourier shell correlation (FSC) plots (left) for the three classes of untreated 30S_Δ*ksg*A_ that refined to resolutions below 5 Å. FSC threshold of 0.143 (dotted line) was used to report the overall resolution of the maps. Cryo-EM density maps (right) colored according to their local resolution. Local resolution scale bar and structural landmarks of the 30S subunit are indicated (top).

**Supplementary Figure 5.**
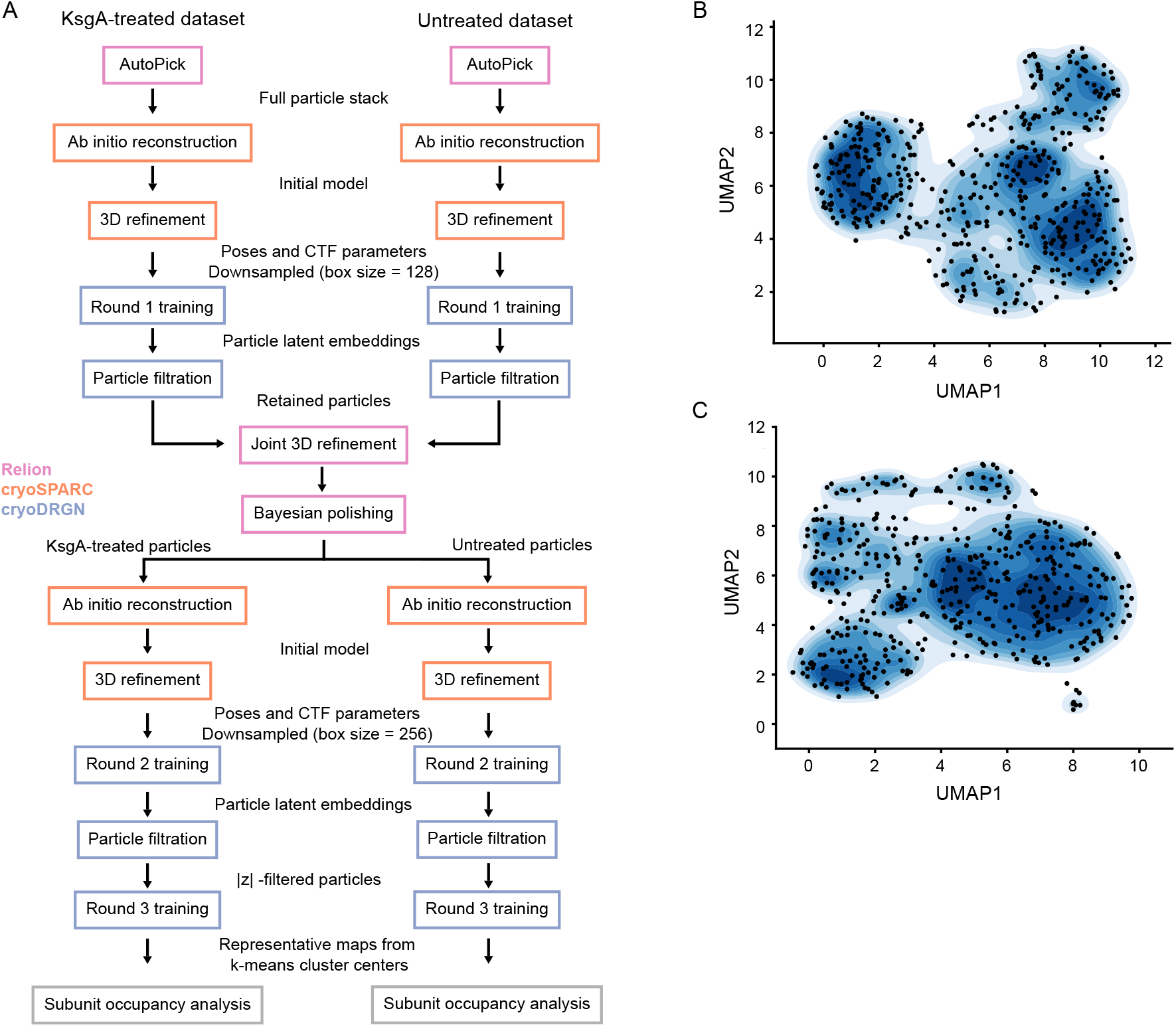
CryoDRGN analysis of untreated and KsgA-treated 30S_Δ*ksg*A_ datasets. **(A)** CryoDRGN data analysis pipeline. Particle filtration following Round 1 training relied on UMAP embedding of the latent space, as described in the Methods. Filtering following Round 2 training used either the magnitude of the eight-dimensional latent z variable, or eliminated whole k-means clusters of particles (see Methods). Additional *ab initio* reconstructions and 3D refinements following Bayesian polishing improved pose assignments on a subset of the particles. UMAP projections of the latent embeddings from the KsgA-treated **(B)** and untreated **(C)** “Round 3” training runs. Projections colored by particle density, and points in latent space sampled for occupancy analysis are noted with black circles.

**Supplementary Figure 6.**
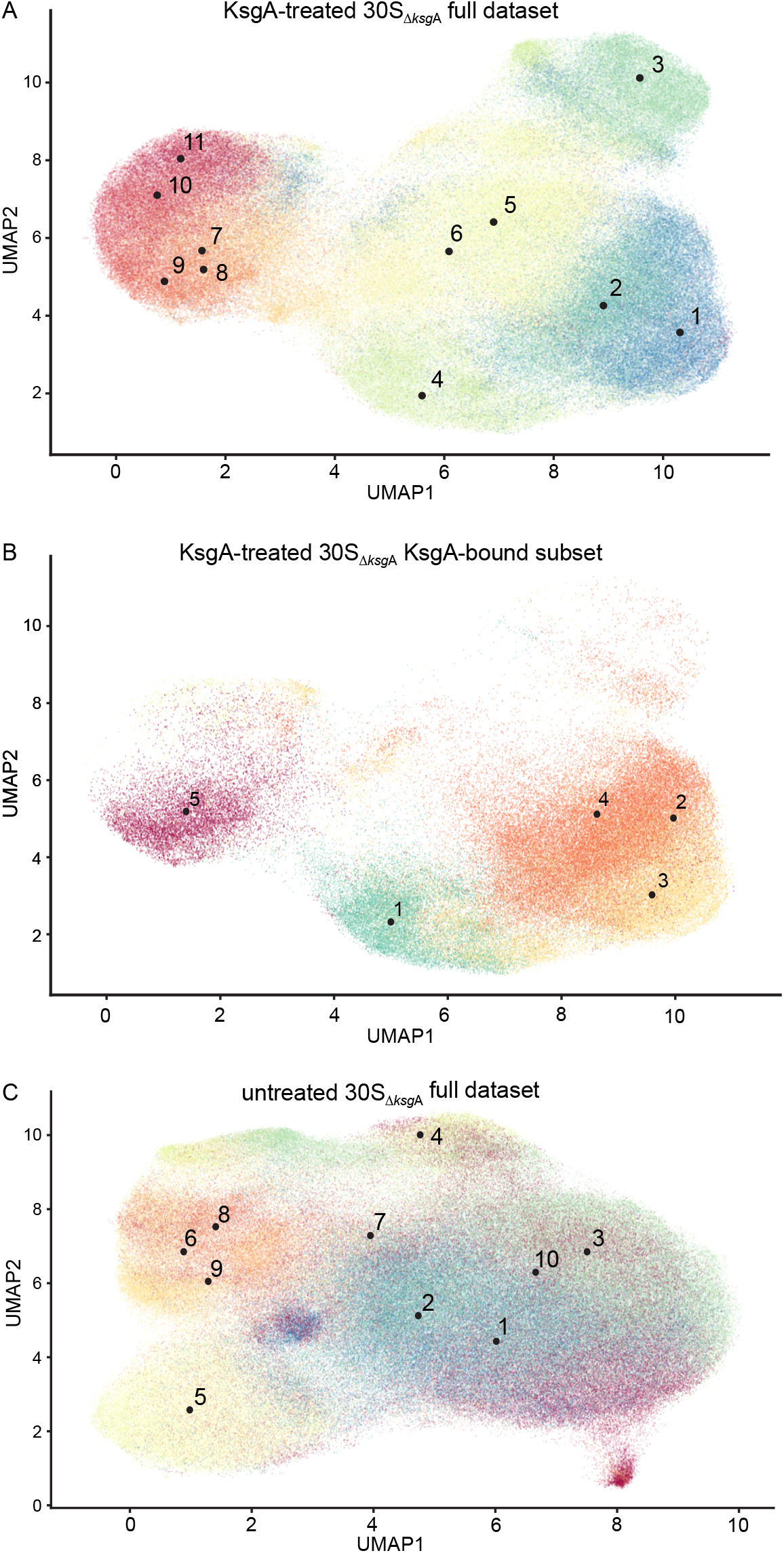
Distribution of cryoDRGN particle embeddings. Two-dimensional UMAP projection of the eight-dimensional latent space, where each point represents the latent embedding of a single particle. Points are colored according to the volume class to which that particle was assigned, as determined by subunit occupancy analysis. Labeled black dots represent the location from which the representative density map was generated for each particle class. Plots depict this analysis for full KsgA-treated 30S_Δ*ksg*A_ dataset **(A)**, the KsgA-bound subset of the KsgA-treated 30S_Δ*ksg*A_ dataset **(B)**, and the full untreated 30S_Δ*ksg*A_ dataset **(C)**.

**Supplementary Figure 7.**
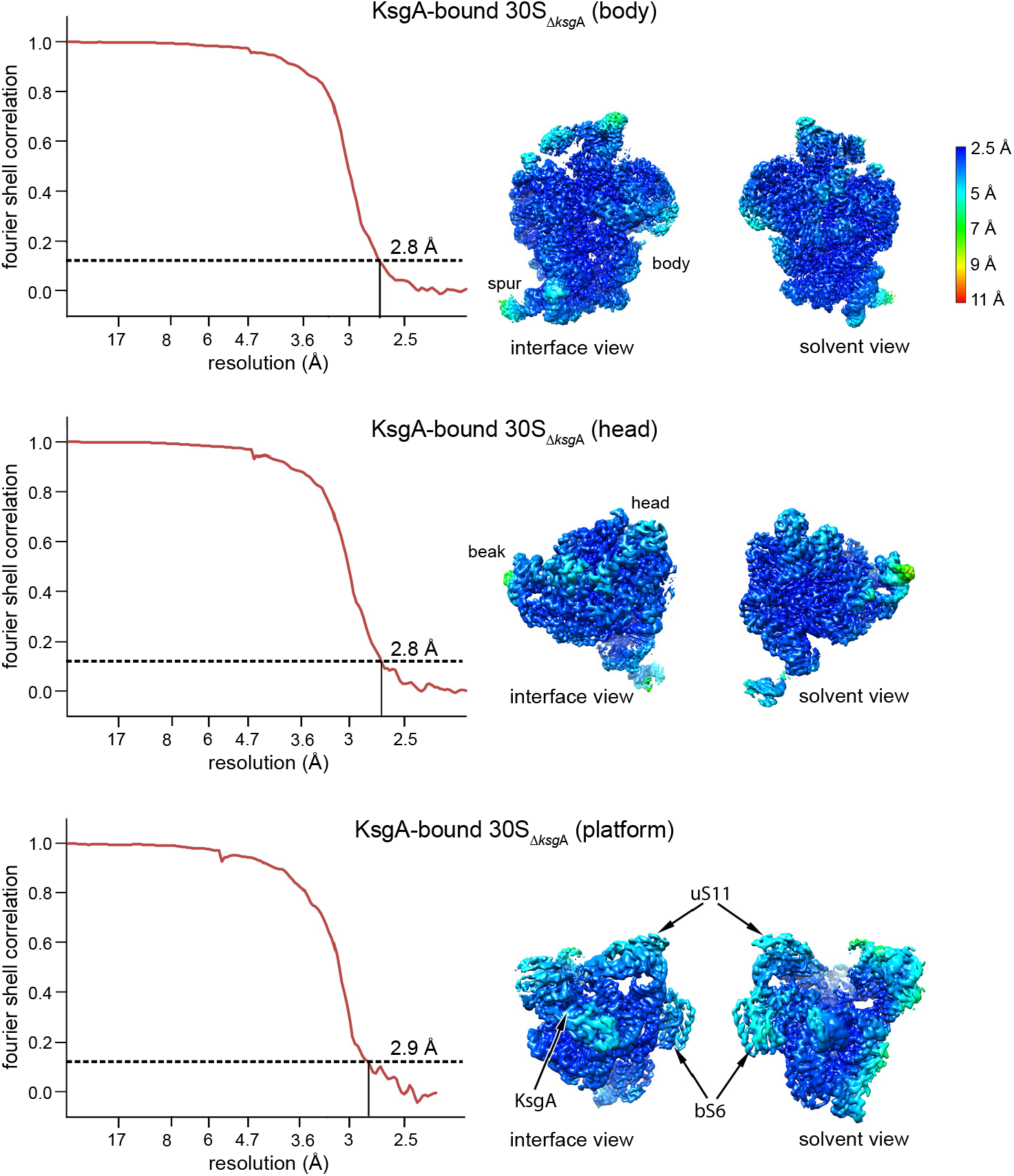
Resolution analysis of the KsgA-bound 30S_Δ*ksg*A_ cryo-EM maps. Fourier shell correlation plots (left) for KsgA-treated 30S_Δ*ksg*A_ maps resulting from multi-body refinement of the body (top), head (middle) and platform (bottom) domains. FSC threshold of 0.143 is used to report the overall resolution of the maps. Cryo-EM maps (right) are colored according to their local resolution using the color coding indicated in the scale bar. Structural landmarks of the 30S subunit are indicated.

**Supplementary Figure 8.**
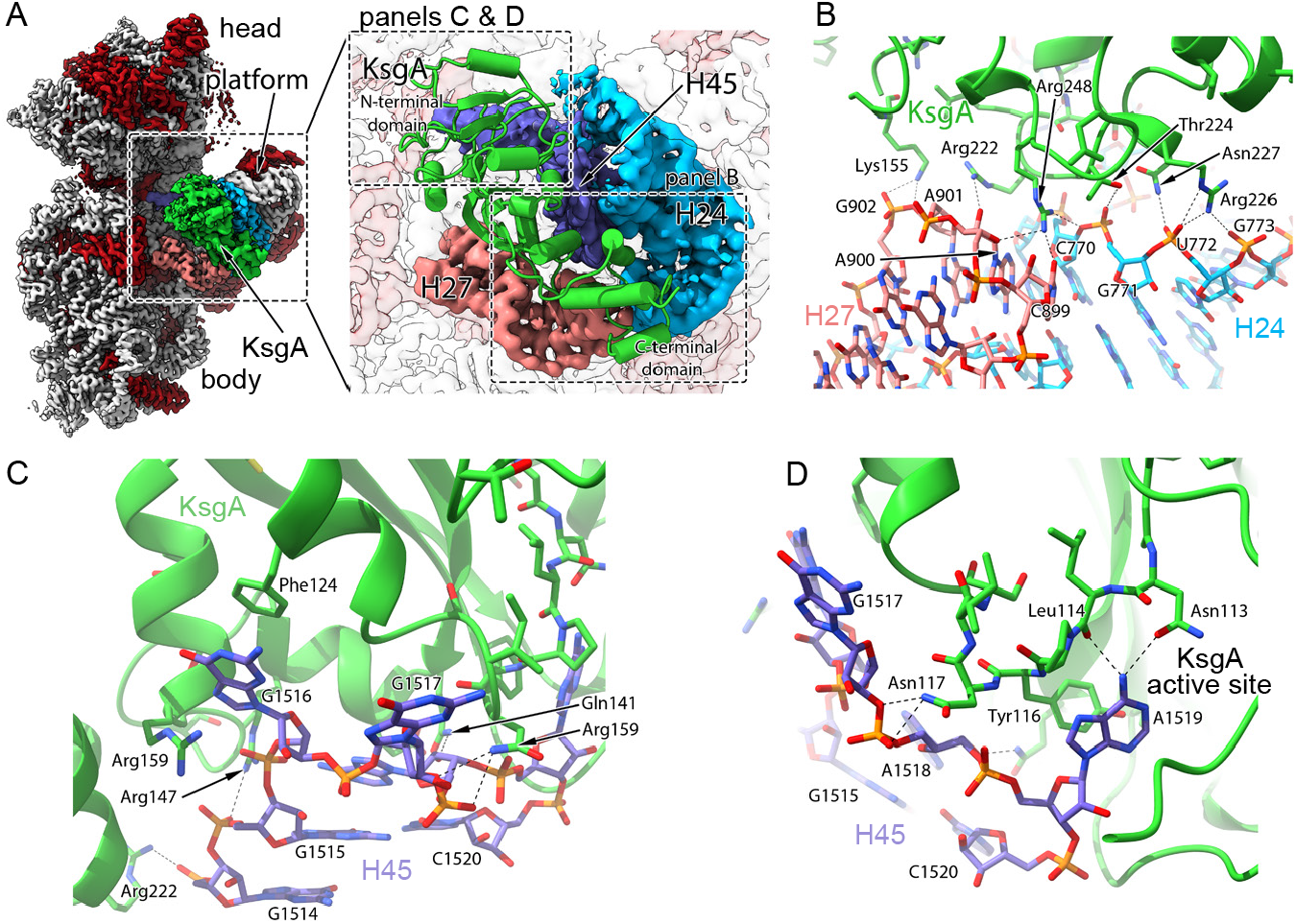
RNA backbone contacts support binding of KsgA. **(A)** Overview of the cryo-EM map obtained for the immature 30S_Δ*ksg*A_ particle bound to KsgA. The framed area is shown magnified in the right panel to highlight the main rRNA helices 24, 27 and 45 that interact with KsgA (green). Views of the molecular model showing the interaction details between KsgA and helices 24, 27 **(B)** and 45 **(C & D)** of the 30S_Δ*ksg*A_ particle.

**Supplemental Figure 9:**
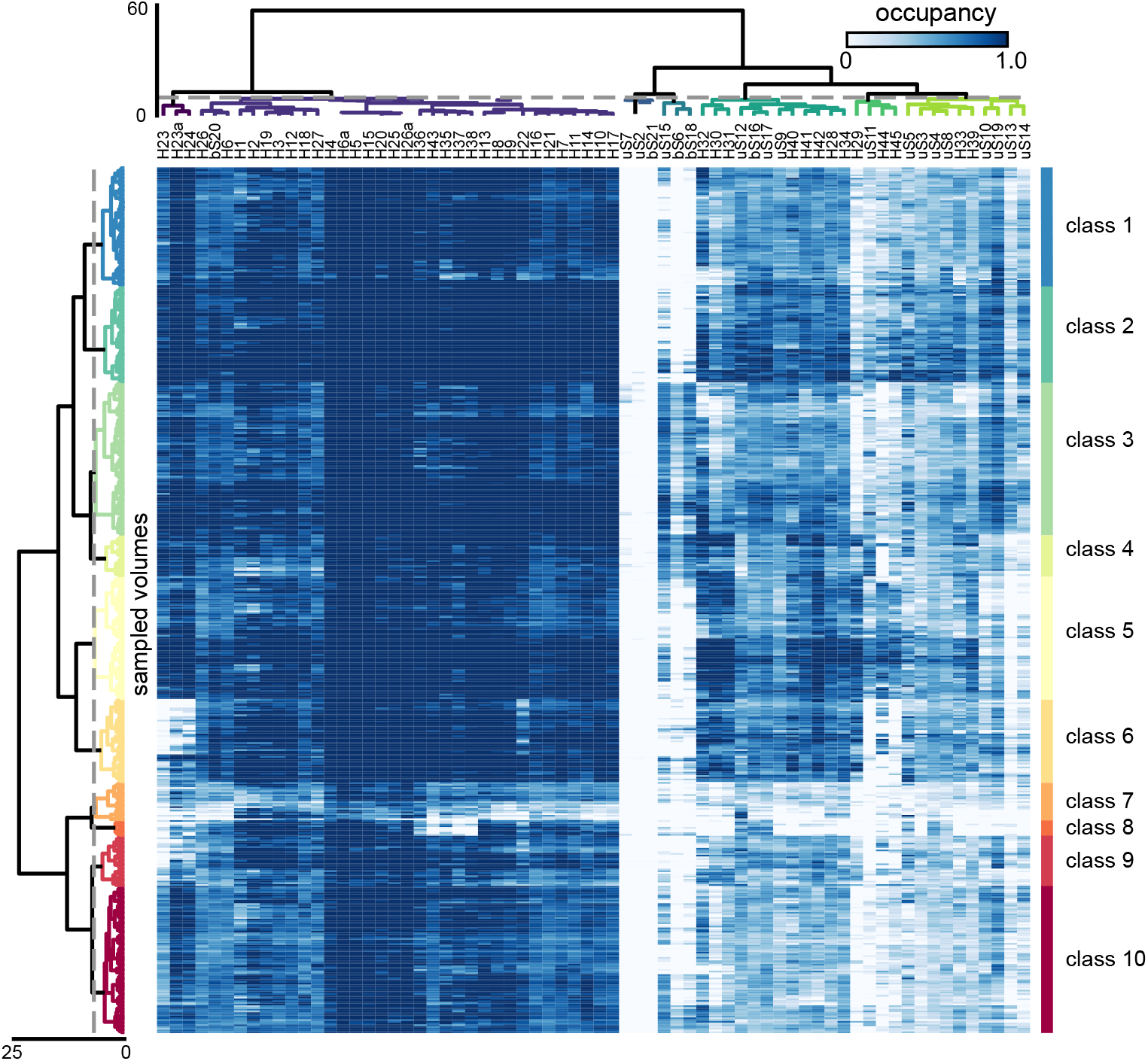
Occupancy analysis of the untreated 30S_Δ*ksg*A_ dataset. A cryoDRGN model was trained on the untreated dataset, and 500 maps were sampled systematically from the resulting latent space. Occupancy analysis was performed as described in the Methods. Rows (500) correspond to sampled maps and columns (68) correspond to structural elements. KsgA occupancy was not measured for this dataset. Dashed gray lines represent the thresholds applied to the row and column dendrograms to define volume classes and structural blocks, respectively.

**Supplementary Figure 10:**
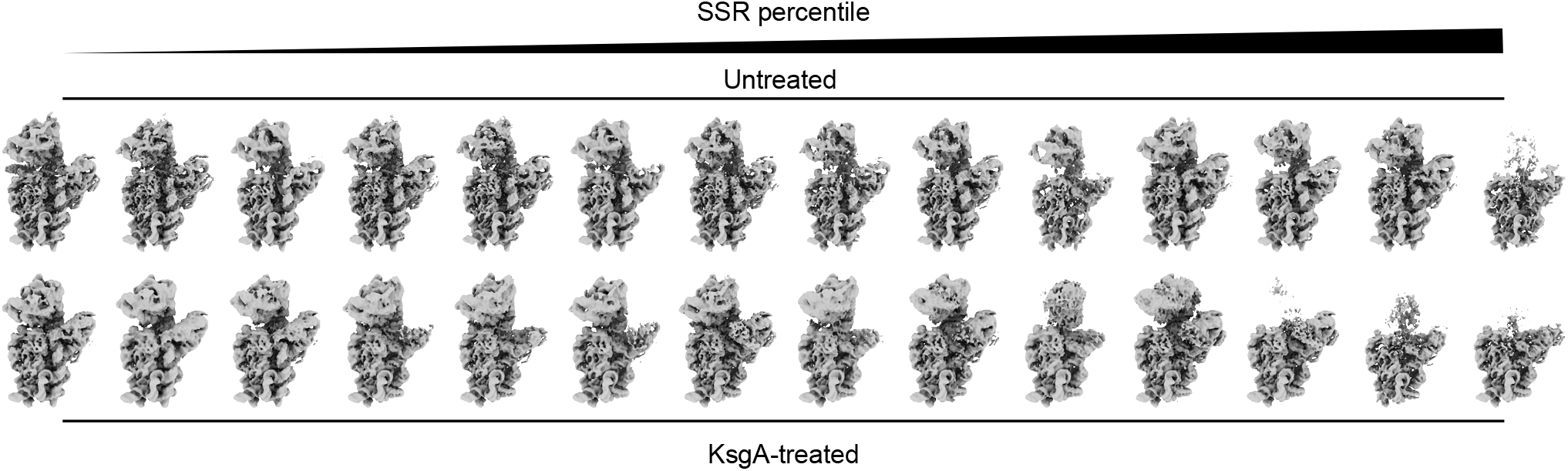
Assessing gross morphology of untreated and KsgA-treated 30S_Δ*ksg*A_ particles. For each of 500 cryoDRGN reconstructed density maps from the untreated and KsgA-treated datasets, a summed squared residual (SSR) value was calculated relative to a mature 30S structure selected from the same dataset (the class 3 centroid volume in each dataset, see Methods). These values were sorted from most-to-least similar to the mature structure, and 14 maps sampled at evenly-spaced percentiles between the 2^nd^ and 98^th^ percentile for each dataset are depicted in ascending order from left to right. The top row shows the percentile maps from the untreated dataset; the bottom row shows the percentile maps from the KsgA-treated dataset

**Supplementary Figure 11.**
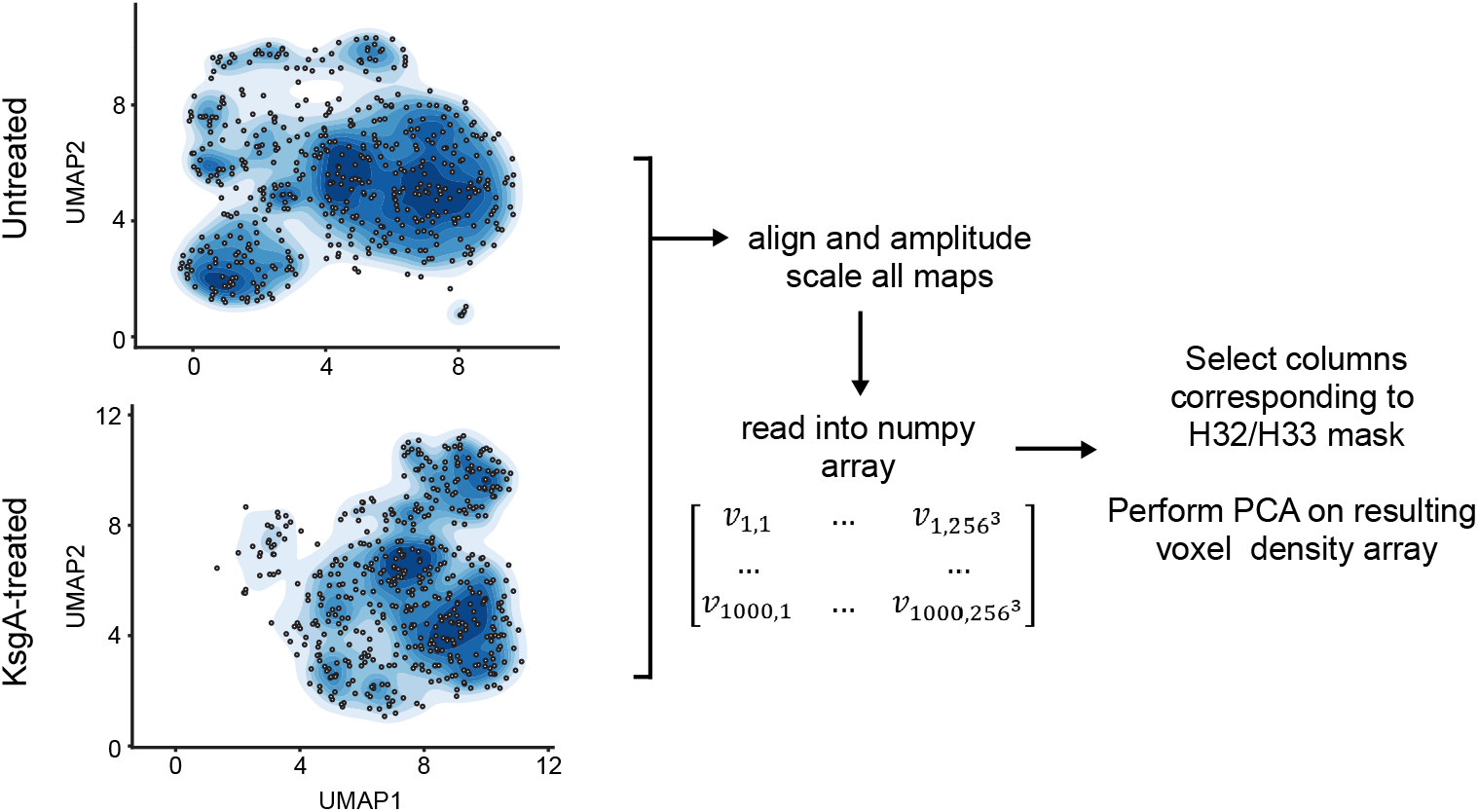
Voxel PCA approach to analyze head domain motions. The subset of particles from each dataset belonging to subunit occupancy analysis classes containing strong head density were identified, and these particle subsets were re-classified by k-means clustering with k = 500. The 1,000 density maps that resulted from cryoDRGN reconstructions from the latent space locations of these k-means cluster centers were aligned and amplitude-scaled (see Methods). A mask corresponding to native H32/33 density was applied to the volumes, and PCA was performed directly on the resulting voxel array.

**Supplementary Figure 12:**
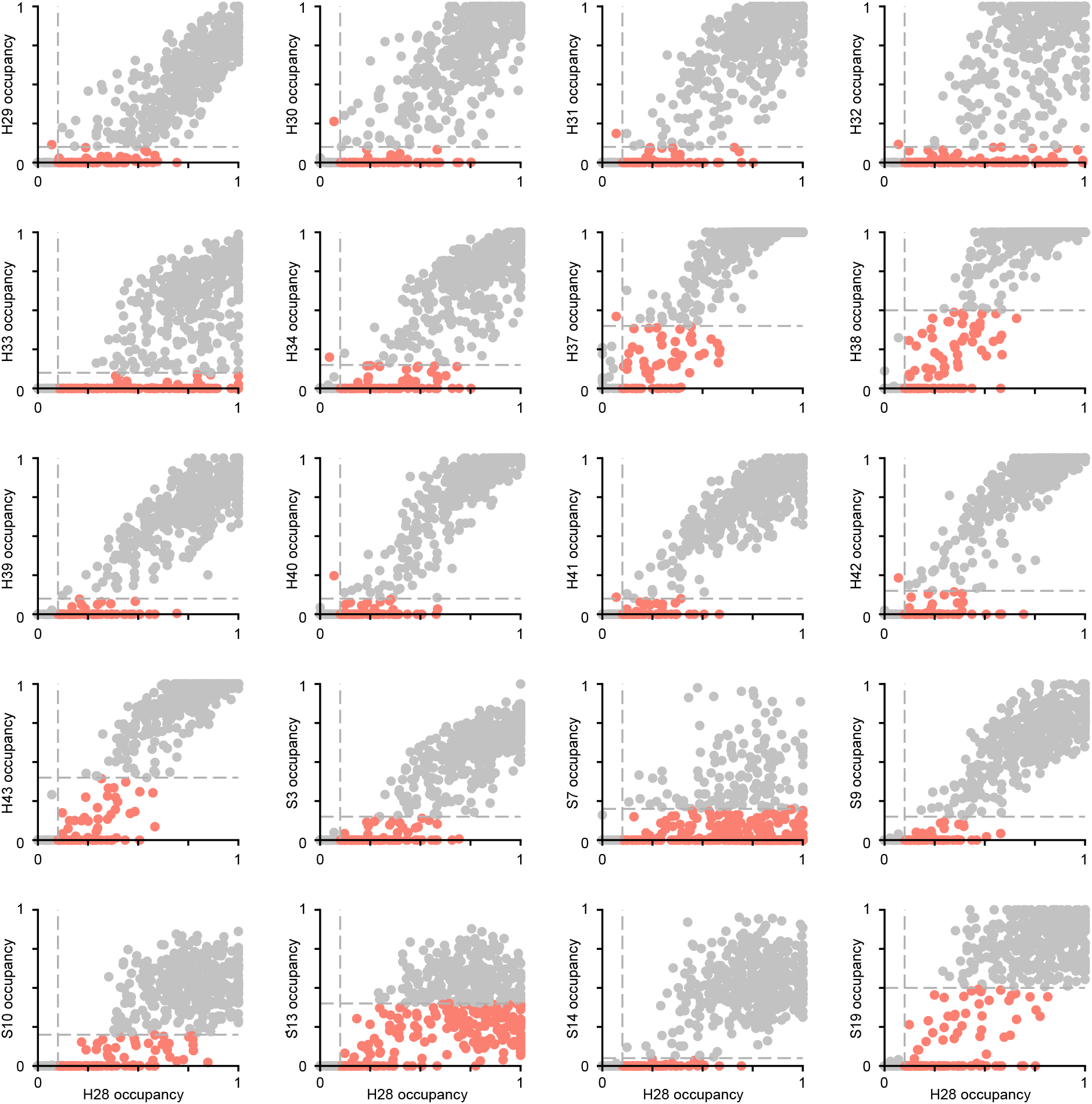
Occupancy of head elements depends on occupancy of H28. Scatter plots comparing occupancy of H28 with each head rRNA helix and r-protein are shown. Each dot represents one volume. Dashed gray lines indicate thresholds dividing high- and low-occupancy volumes for each subunit (see Methods). Dots are colored to indicate dependency – volumes with low occupancy of one subunit and high occupancy of the other are colored red, while volumes that are not informative for dependency (both highly-occupied or both lowly-occupied) are colored gray.

**Supplementary Video 1: Rotation of the head density in KsgA-treated volumes**.

For each of the KsgA-treated and untreated datasets, 500 volumes were resampled from the subset of latent space corresponding to particles that belonged to classes exhibiting strong head density. The full set of 1,000 volumes were aligned and amplitude-scaled, and a mask corresponding to the native H32/H33 density in PDB model 4V9D was applied to the volumes. Principal component analysis was performed on the resulting voxel array (see Methods and Supplemental Figure 11). Volumes shown here are from the KsgA-treated dataset, sampled from low to high values of the first principal component. Only the head portion of each volume is shown (orange); the mature 30S is overlaid for reference (grey).

**Supplementary Video 2: H44 flexibility in the absence of KsgA**.

For each of the treated and untreated datasets, 500 volumes were resampled from the subset of latent space corresponding to particles that belonged to classes exhibiting strong H44 density. The full set of 1,000 volumes were aligned and amplitude-scaled, and a mask was applied to the volumes, corresponding to a 12 Å-expanded region around H44 in PDB model 4V9D. Principal component analysis was performed on the resulting voxel array (see Methods and Supplemental Figure 11). Volumes shown here are from the untreated dataset, sampled from low to high value of the first principal component. Only the H44-masked portion of each volume is shown (orange); a volume lacking H44 is overlaid for reference (grey).

**Supplementary Figure 13:**
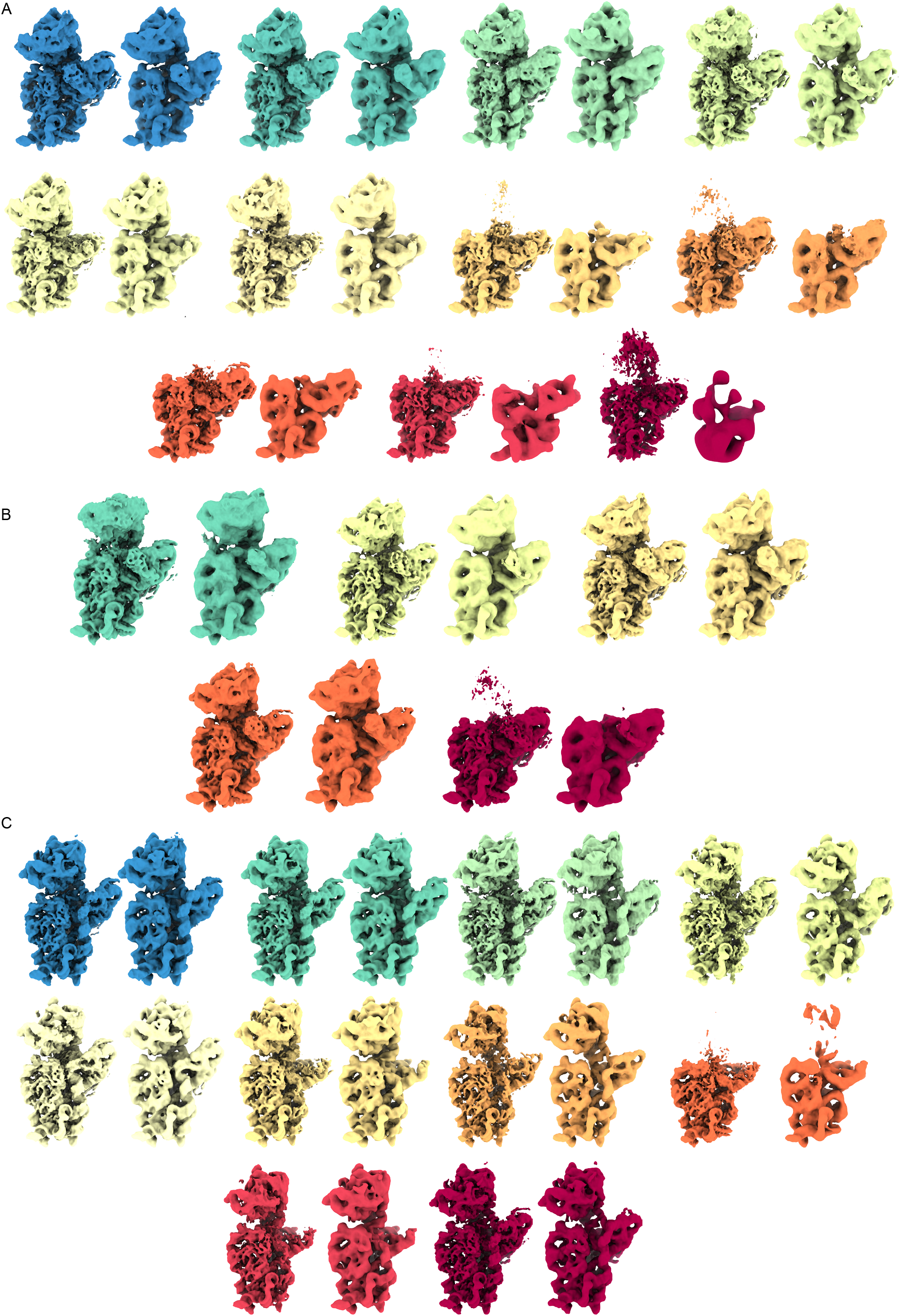
Validation of centroid volumes via traditional processing workflows. For each class centroid volume identified via subunit occupancy analysis, the nearest 1% of particles in the corresponding latent space were identified, and *ab initio* model generation and homogeneous refinement were carried out in cryoSPARC using these particles. Class centroid volumes (left volume of each colored pair) and corresponding homogeneous refinements (right volume of each colored pair) are shown for **(A)** the KsgA-treated dataset, with all volumes clustered (see Figure 2), **(B)** the KsgA-treated dataset, with only volumes having a KsgA occupancy > 0.15 clustered (see Figure 3), and C) the untreated dataset (see Figure 5). Volumes are colored to match the corresponding classes and volumes in the relevant main-text figures.

